# Interdependence of plasma membrane nanoscale dynamics of a kinase and its cognate substrate underlies *Arabidopsis* response to viral infection

**DOI:** 10.1101/2023.07.31.551174

**Authors:** Marie-Dominique Jolivet, Anne-Flore Deroubaix, Marie Boudsocq, Nikolaj B. Abel, Marion Rocher, Terezinha Robbe, Valérie Wattelet-Boyer, Jennifer Huard, Dorian Lefebvre, Yi-Ju Lu, Brad Day, Grégoire Saias, Jahed Ahmed, Valérie Cotelle, Nathalie Giovinazzo, Jean-Luc Gallois, Yasuyuki Yamaji, Sylvie German-Retana, Julien Gronnier, Thomas Ott, Sébastien Mongrand, Véronique Germain

**Author notes:** Corresponding author: Véronique Germain. Nikolaj B. Abel: Aarhus University, 8000 Aarhus, Denmark.

## Abstract

Plant viruses represent a risk to agricultural production and as only a few treatments exist, it is urgent to identify resistance mechanisms and factors. In plant immunity, plasma membrane (PM)-localized proteins play an essential role in sensing the extracellular threat presented by bacteria, fungi or herbivores. Viruses are intracellular pathogens and as such the role of the plant PM in detection and resistance against viruses is often overlooked. We investigated the role of the partially PM-bound Calcium-dependent protein kinase 3 (CPK3) in viral infection and we discovered that it displayed a specific ability to hamper viral propagation over CPK isoforms that are involved in immune response to extracellular pathogens. More and more evidence support that the lateral organization of PM proteins and lipids underlies signal transduction in plants. We showed here that CPK3 diffusion in the PM is reduced upon activation as well as upon viral infection and that such immobilization depended on its substrate, Remorin (REM1.2), a scaffold protein. Furthermore, we discovered that the viral infection induced a CPK3-dependent increase of REM1.2 PM diffusion. Such interdependence was also observable regarding viral propagation. This study unveils a complex relationship between a kinase and its substrate that contrasts with the commonly described co-stabilisation upon activation while it proposes a PM-based mechanism involved in decreased sensitivity to viral infection in plants.

## INTRODUCTION

Viruses are intracellular pathogens, carrying minimal biological material and strictly relying on their host for replication and propagation. They represent a critical threat to both human health and food security. In particular, potexvirus epidemics like the one caused by pepino mosaic virus dramatically affect crop production^1^ and the lack of chemical treatments available makes it crucial to develop inventive protective methods. Unlike their animal counterparts, which enter host cells by interacting directly with the plasma membrane (PM), plant viruses have to rely either on mechanical wounding or insect vectors to cross the plant cell wall^1^. For this reason, only a few PM-localized proteins were identified as taking part in immunity against viruses^2,3^. Among them, members of the REMORIN (REM) protein family were shown to be involved in viral propagation, with varying mechanisms depending on the studied viral genera^4–9^. REM proteins are well-known for their heterogeneous distribution at the PM, forming nanodomains (ND), PM nanoscale environments that display a composition different from the surrounding PM^10,11^. Increasing evidence supports the role of ND in signal transduction, with the underlying idea that the local accumulation of proteins allows amplification and specification of the signal^11^. For example, *Arabidopsis* RHO-OF-PLANT 6 accumulates in distinct ND upon osmotic stress and auxin treatment in a dose-dependent way for the latter^12^. Recently, *Arabidopsis* REM1.2 (later named REM1.2) was demonstrated to form clusters upon exposure to the bacterial effector flg22 to support the condensation of *Arabidopsis* FORMIN 6 and to induce actin cytoskeleton remodeling^13^. However, unlike the canonical mechanism describing the accumulation of proteins in ND upon stimulation^12–14^, we showed previously that *Solanum tuberosum* REM1.3 (StREM1.3) ND were disrupted and the diffusion of individual proteins increased in response to a viral infection^15^. StREM1.3 lateral organization in lipid bilayers was also shown to be modified upon its phosphorylation status, both *in vitro* and *in vivo*^15,16^. The role of such protein dispersion upon stimuli is not understood yet and only a few similar cases are reported in the literature^17,18^. REM1.2 was identified as a substrate of the partially membrane-bound CALCIUM-DEPENDENT PROTEIN KINASE 3 (CPK3)^19^. CPK3 is involved in defense response against herbivores, bacteria and viruses and was recently proposed to be at cross-roads between pattern-triggered immunity and effector-triggered immunity^15,20,21^. We showed previously that transient over-expression of CPK3 in *N. benthamiana* was able to hamper potato virus X (PVX, potexvirus) cell-to-cell propagation^15^. Although partially PM localized, the role of CPK3 PM localization in immunity was never investigated for any pathogen. As the PVX cannot infect *Arabidopsis thaliana*, the dedicated plant model for genetic studies, we used an alternative virus model able to infect this species, the plantago asiatica mosaic virus (PlAMV)^22,23^ to investigate the role of CPK3 and of its PM localization in potexvirus propagation. We were able to highlight the specific role of CPK3 among other immune-related CPKs^24–26^ in this context. Also, we demonstrated that CPK3 membrane localization was required to hamper viral cell-to-cell propagation and, using single-particle tracking photoactivated light microscopy (spt-PALM), we discovered that CPK3 diffusion was reduced upon activation and viral infection. Interestingly, this reduction of PM diffusion depended on the expression of Group 1 REM while viral-induced REM1.2 increased PM diffusion depended on CPK3. Overall, our data allowed us to propose a model for a PM-localized mechanism involved in the reduction of potexvirus propagation, which will encourage further exploration of the involvement of the PM in viral immunity.

## RESULTS

### *Arabidopsis thaliana* calcium-dependent protein kinase 3 (CPK3) specifically restricts PlAMV propagation

We previously showed that transient overexpression of *Arabidopsis* CPK3 in *N. benthamiana* leaves restricted the propagation of the potexvirus potato virus X (PVX)^15^. CPKs are encoded by a large gene family of 34 members in *Arabidopsis*^27^. To evaluate the functional redundancy between CPKs regarding viral propagation, a series of *Arabidopsis* lines mutated for CPK1, CPK2, CPK3, CPK5, CPK6 or CPK11, that are involved in plant resistance to various pathogens^21,24–26,28^, were analyzed in a viral propagation assay. Because PVX does not infect *Arabidopsis*, we used instead a binary plasmid encoding for the genomic structure of a GFP-tagged PlAMV^22,23^, a potexvirus that is capable of infecting a wide range of plant hosts, including *Arabidopsis*. Agrobacterium carrying PlAMV-GFP were infiltrated in *A. thaliana* leaves and GFP-fluorescent infection foci were observed 5 days post infiltration (dpi) (Figure 1 – figure supplement 1). The following combinations of mutants were tested: the *cpk1 cpk2* double mutant^25^, the *cpk5 cpk6* double mutant^24^, the triple mutant *cpk5 cpk6 cpk11*^24^ and the two quadruple mutants *cpk1 cpk2 cpk5 cpk6*^25^ and *cpk3 cpk5 cpk6 cpk11*^26^. No significant difference of PlAMV-GFP infection foci area was detected between Col-0, *cpk1 cpk2, cpk5 cpk6, cpk1 cpk2 cpk5 cpk6* and *cpk5 cpk6 cpk11,* demonstrating that CPK1, CPK2, CPK5, CPK6 and CPK11 are not involved in PlAMV propagation (Figure 1A, 1B). However, a 20 % increase of PlAMV-GFP infected area was observed in *cpk3 cpk5 cpk6 cpk11* quadruple mutant relative to the control Col-0, which indicates that CPK3 could play a specific role in viral propagation. Since group 1 REMs are known substrates of CPK3^19^, we checked REM1.2 phosphorylation specificity by the CPK isoforms tested in viral propagation (Figure 1 – figure supplement 2). CPK1, 2 and 3 displayed the strongest kinase activity on REM1.2 while CPK5, CPK6 and CPK11 displayed a residual activity. In contrast, all 6 CPKs phosphorylated the generic substrate histone, suggesting some substrate specificity for REM1.2 *in vitro*. Since CPK1 and CPK2 were described to be mainly localized within the peroxisome^29^ and endoplasmic reticulum^30^, respectively, we hypothesize that they likely do not interact in *vivo* with PM-localized REM1.2^31^.

**Figure 1.**
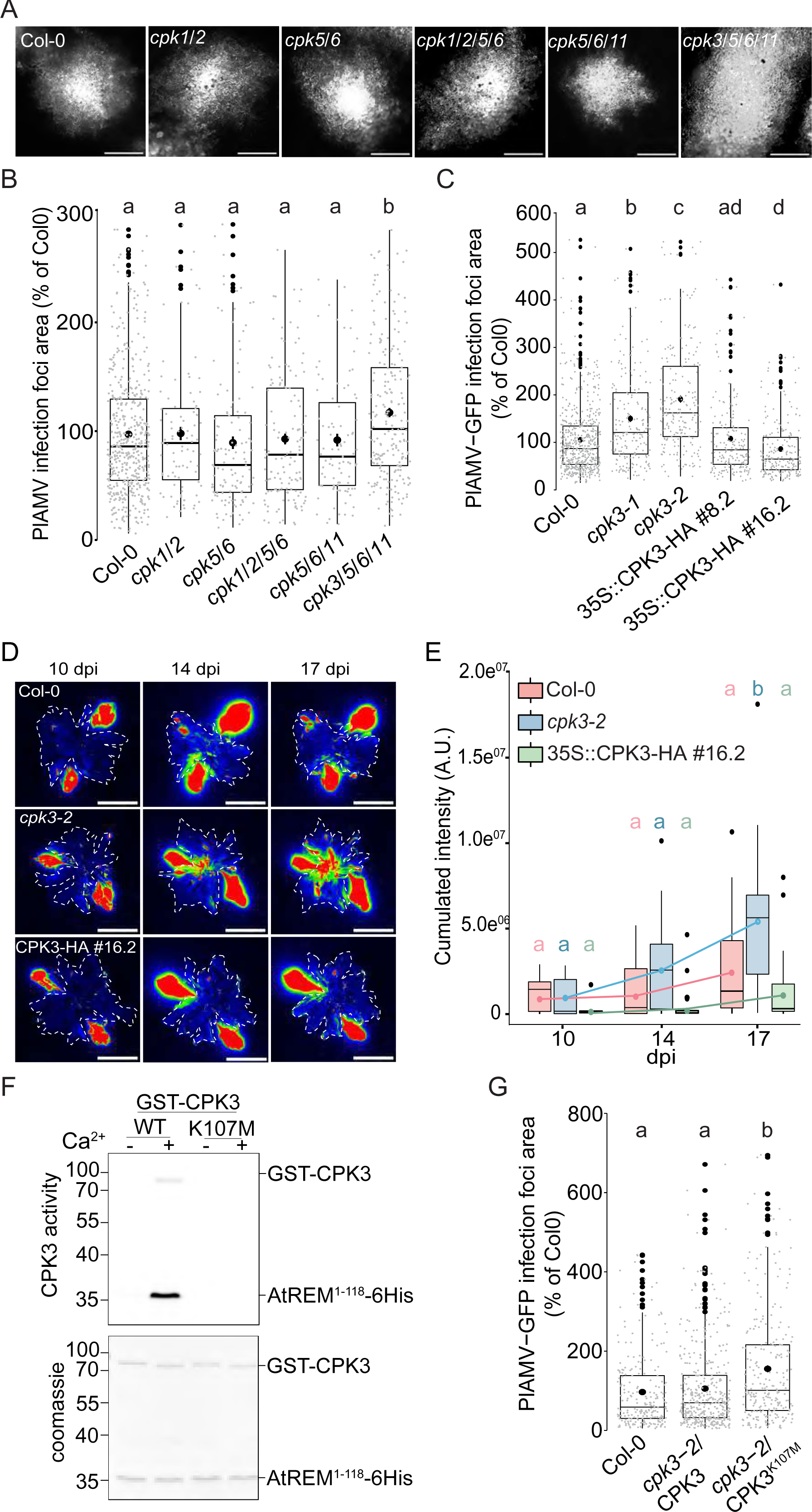
*Arabidopsis thaliana* calcium-dependent protein kinase 3 (CPK3) is specifically involved in the restriction of PlAMV cell-to-cell movement. A. Representative images of PlAMV-GFP infection foci at 5 dpi in the different mutant backgrounds. Scale bar = 500 µm B. Box plots of the mean area of PlAMV-GFP infection foci 5 days after infection in CPK multiple mutant lines, normalized to the mean area measured in Col-0. Three independent biological repeats were performed, with at least 47 foci per experiment and per genotype. Significant differences were revealed using a One-Way ANOVA followed by a Tukey’s multiple comparison test. Letters are used to discriminate between statistically different conditions (p<0.05). C. Box plots of the mean area of PlAMV-GFP infection foci in *cpk3-1* and *cpk3-2* single mutants and in CPK3 over-expressing lines (Pro35S:CPK3-HA #8.2 and Pro35S:CPK3-HA #16.2), normalized to the mean area measured in Col-0. Three independent biological repeats were performed, with at least 56 foci per experiment and per genotype. Significant differences were revealed using a One-Way ANOVA followed by a Tukey’s multiple comparison test. Letters are used to discriminate between statistically different conditions (p<0.05). D. Representative images of *A. thaliana* plants infected with PlAMV-GFP and imaged with a CCD Camera from 10 to 17 dpi. The region of interest used for measurement of pixel intensity is circled with a white dotted line. Multicolored scale is used to enhance contrast and ranges from blue (low intensity) to red (high intensity). Scale bar = 4 cm. E. Box plots of the mean cumulated intensity measured in infected leaves in Col-0, *cpk3-2* and Pro35S:CPK3-HA #16.2 during systemic viral propagation. Two independent experiments were conducted. Statistical differences could be observed between the genotypes and time-points using a Kruskal-Wallis followed by a Dunn’s multiple comparison test (p<0.05). For clarity, only the results of the statistical test of the comparison of the different time-points (10, 14 and 17 dpi) within a genotype are displayed and are color-coded depending on the genotype. F. Kinase activity of CPK3 dead variant. Recombinant proteins GST-CPK3 WT and K107M were incubated with REM1.2^1–118^–6His in kinase reaction buffer in the presence of EGTA (−) or 100 µM free Ca^2+^ (+). Radioactivity is detected on dried gel (upper panel). The protein amount is monitored by Coomassie staining (lower panel). G. Box plots of the mean area of PlAMV-GFP infection foci in *cpk3-2* complemented lines *cpk3-2*/Pro35S:CPK3-myc and *cpk3-2*/ Pro35S:CPK3^K107M^-myc. Three independent biological repeats were performed, with at least 51 foci per experiment and per genotype. Significant differences were revealed using a One-Way ANOVA followed by a Tukey’s multiple comparison test. Letters are used to discriminate between statistically different conditions (p<0.0001)

To further confirm a role for CPK3 in PlAMV infection, two independent knock-out (KO) lines for CPK3, *cpk3-1*^32^ and *cpk3-2*^19^ along with two independent lines overexpressing CPK3 (i.e., Pro35S:CPK3-HA-OE #8.2 and Pro35S:CPK3-HA-OE#16.2^72^) (Figure 1 – figure supplement 3) were infiltrated with PlAMV-GFP. In both *cpk3-1* and *cpk3-2* KO lines, PlAMV-GFP propagation was significantly enhanced (40 to 60% compared with WT Col-0), whereas the over-expression lines Pro35S:CPK3-HA-OE#8.2 and Pro35S:CPK3-HA-OE#16.272 showed 10% and 20 % restriction of the foci area, respectively (Figure 1C). To assess whether CPK3 regulates viral propagation at the plant level, the propagation of PlAMV-GFP in systemic leaves was assessed in 3-week-old *cpk3-2* and Pro35S:CPK3-HA-OE#16.2 lines along with Col-0 at 10, 14 and 17 days post infection (dpi) using a CCD Camera equipped with a GFP filter (Figure 1 – figure supplement 1). We observed that loss of CPK3 led to an increase of PlAMV-GFP propagation in distal leaves during the course of our assay while the overexpression of CPK3 did not hamper PlAMV-GFP to a greater extent than WT Col-0 (Figure 1D and E). This would suggest that CPK3 effect on PlAMV propagation is predominant at the site of infection. For this reason, we concentrated on foci in local leaves for further study.

CPK3 Lysine 107 functions as an ATP binding site and its substitution into methionine abolishes CPK3 activity *in vitro*^21^. In good agreement, we observed that CPK3^K107M^ can no longer phosphorylate REM1.2 *in vitro* (Figure 1F). To test whether CPK3 kinase activity is required for its function during PlAMV infection, we analyzed the propagation of PlAMV-GFP in complementation lines of *cpk3-2* with WT CPK3 or with CPK3^K107M^. While the WT CPK3 fully complemented *cpk3-2* mutant, it was not the case with *cpk3-2*/Pro35S:CPK3^K107M^, which displayed larger infection foci area compared to WT Col-0 (Figure 1G).

Taken together, these results demonstrate a specific involvement for CPK3 among other previously described immune-related CPKs in limiting PlAMV infection.

### PlAMV infection induces a decrease in CPK3 PM diffusion

We next wondered whether CPK3 accumulation is regulated at transcriptional and translational level upon PlAMV infection. RT-qPCR and western blots of CPK3 in Col-0 showed that both transcript and protein levels remained unchanged during PlAMV infection (Figure 2 – figure supplement 1). CPK3 is partially localized within the PM and is myristoylated at Glycine 2, a modification required for its association with membranes^19^. To test whether CPK3 membrane localization is required to hamper PlAMV propagation, we transformed *cpk3-2* mutant with either ProUbi10:CPK3-mRFP1.2 or ProUbi10:CPK3^G2A^-mRFP1.2 (Figure 2A) and tested PlAMV infection. We observed that, in contrary to ProUbi10:CPK3-mRFP1.2, ProUbi10:CPK3^G2A^-mRFP1.2 did not complement *cpk3-2* (Figure 2B). These observations indicate that CPK3 association with the PM is required for its function in inhibiting PlAMV propagation. We next analyzed the organization of CPK3 PM pool in absence or presence of PlAMV-GFP using confocal microscopy. Imaging of the surface of *A. thaliana* leaf epidermal cells expressing *CPK3-mRFP1.2* showed that the protein displayed a heterogeneous pattern at the PM in both conditions (Figure 2C), although the limitation in lateral resolution of confocal microscopy hindered a more detailed analysis of CPK3 PM organization.

**Figure 2.**
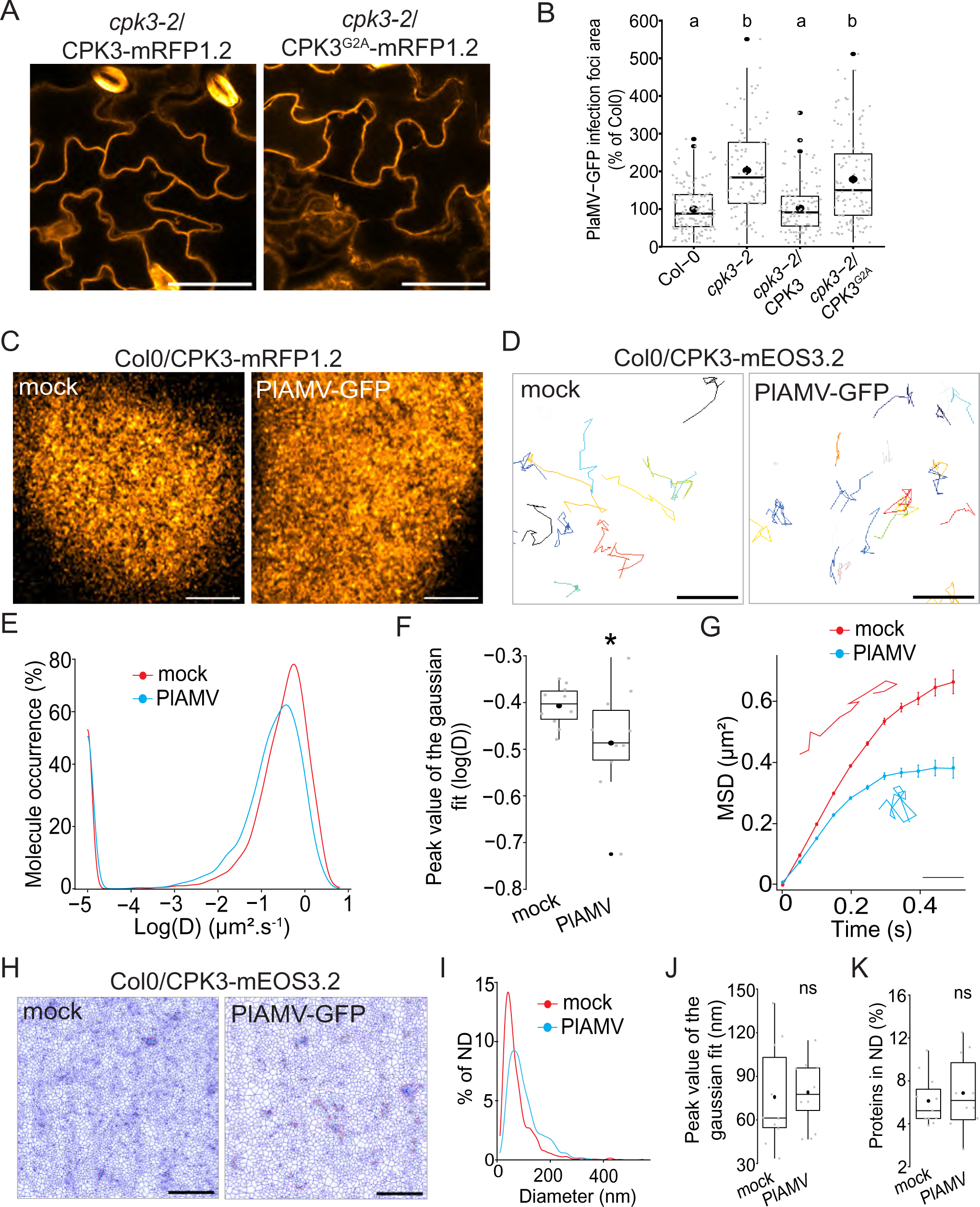
CPK3 diffusion decreases upon PlAMV infection in *Arabidopsis thaliana*. A. Confocal images of the secant view of *A. thaliana* epidermal cells of the *cpk3-2*/CPK3-mRFP1.2 or *cpk3-2*/ProUbi10:CPK3^G2A^-mRFP1.2. Scale bar = 5µm B. Box plots of the mean area of PlAMV-GFP infection foci in *cpk3-2* and complemented lines *cpk3-2*/ProUbi10:CPK3-mRFP1.2 and *cpk3-2*/ProUbi10:CPK3^G2A^-mRFP1.2. Three independent biological repeats were performed, with at least 32 foci per experiment and per genotype. Significant differences were revealed using a One-Way ANOVA followed by a Tukey’s multiple comparison test. Letters are used to discriminate between statistically different conditions (p<0.0001). C. Confocal images of the surface view of *A. thaliana* epidermal cells of the Col-0/ProUbi10:CPK3-mRFP1.2, infiltrated either with free GFP (“mock”) or PlAMV-GFP. Scale bar = 5µm D. Representative trajectories of Col-0/CPK3-mEOS3.2 infiltrated either with free GFP (“mock”) or PlAMV-GFP; Scale bar = 2µm E. Distribution of the diffusion coefficient (D), represented as log(D) for Col-0/ProUbi10:CPK3-mEOS3.2 five days after infiltration with free GFP (“mock”) or PlAMV-GFP. Data were acquired from at least 8086 trajectories obtained in at least 16 cells over the course of three independent experiments. F. Box plot of the mean peak value extracted from the Gaussian fit of log(D) distribution. Significant difference was revealed using a Mann-Whitney test. *: p<0.05. G. Mean square displacement (MSD) over time of Col-0/ProUbi10:CPK3-mEOS3.2 five days after infiltration with free GFP (“mock”) or PlAMV-GFP. Representative trajectories extracted from Figure 2D illustrate each curve. Scale bar = 1µm. Data were acquired from at least 16 cells over the course of three independent experiments. H. Voronoi tessellation illustration of Col-0/ProUbi10:CPK3-mEOS3.2 five days after infiltration with free GFP (“mock”) or PlAMV-GFP. ND are circled in red. Scale bar = 2µm I. Distribution of the ND diameter of Col-0/ProUbi10:CPK3-mEOS3.2 five days after infiltration with free GFP (“mock”) or PlAMV-GFP. J. Box plot representing the mean peak value of ND diameter extracted from the Gaussian fit of Figure 2I. No significant differences were revealed using a Mann-Whitney test. K. Boxplot of the proportion of Col-0/ProUbi10:CPK3-mEOS3.2 detections found in ND five days after infiltration with free GFP (“mock”) or PlAMV-GFP. No significant differences were revealed using a Mann-Whitney test.

Thus, we used single particle tracking phospho-activated localization microscopy (sptPALM) which overcomes the diffraction limit of confocal microscopy and allows the analysis of the diffusion and organization of single molecules. We used a translational fusion of CPK3 with the true monomeric photoconvertible fluorescent protein mEOS3.2^33^ expressed in stable transgenic *Arabidopsis* lines. We imaged these materials in control and upon PlAMV infection. We tracked single molecule trajectories (Figure 2D) from which CPK3 diffusion coefficient (D) was calculated. We observed that CPK3 proteins were overall mobile in control and infected conditions (log(D) > −2; Figure 2E) although CPK3 diffusion was reduced upon PlAMV infection (Figure 2F). Analysis of the mean squared displacement (MSD), describing the surface explored by single molecules overtime, showed that CPK3 displayed a more confined behavior during a PlAMV infection than in healthy conditions (Figure 2G). Additionally, we performed cluster analysis using Voronoï tessellation, a computation method that segments super-resolution images into polygons based on the local molecule density^34^. Voronoï analysis showed that no difference occurred in CPK3 cluster size or proportion of protein localized in cluster upon viral infection (Figure 2H-K). Taken together, these results show that CPK3 diffusion parameters were modified upon PlAMV infection, although the nano-organization of the proteins was maintained.

### Truncation of CPK3 auto-inhibitory domain induces its confinement and accumulation in PM ND

CPK3 bears an auto-inhibitory domain that folds over the kinase domain and inhibits its kinase activity in the absence of calcium^35,36^. The truncation of this domain along with the C-terminal regulatory domain results in a calcium-independent, constitutively active CPK3 (CPK3^CA^)^24^ that is lethal when stably expressed in *Arabidopsis*^37^. For this reason, CPK3^CA^ was transiently expressed in *N. benthamiana* for further analysis. We observed that although both ProUbi10:CPK3-mRFP1.2 and ProUbi10:CPK3^CA^-mRFP1.2 were partially cytosolic when transiently expressed in *N. benthamiana* (Figure 3 – figure supplement 1), ProUbi10:CPK3^CA^-mRFP1.2 displayed a PM organization in domains discernable by confocal microscopy (Figure 3A).

**Figure 3.**
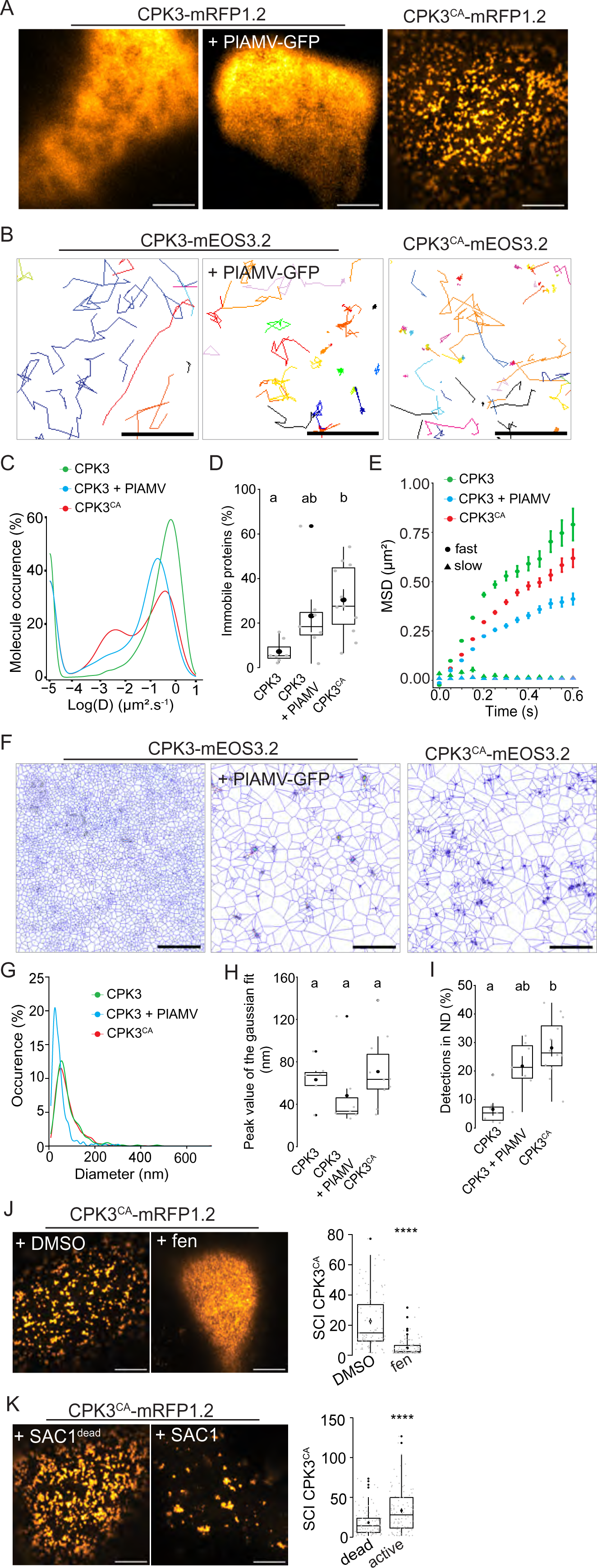
PlAMV-induced activation of CPK3 in *N. benthamiana* induces its confinement and clustering in PM domains. A. Confocal images of the surface view of *N. benthamiana* epidermal cells transiently expressing ProUbi10:CPK3-mRFP1.2, ProUbi10:CPK3-mRFP1.2 + PlAMV-GFP or ProUbi10:CPK3^CA^-mRFP1.2. Scale bar = 5µm B. Representative trajectories of ProUbi10:CPK3-mEOS3.2, ProUbi10:CPK3.2-mEOS3.2 + PlAMV and ProUbi10:CPK3^CA^-mEOS3.2. Scale bar = 2µm. C. Distribution of the diffusion coefficient (D), represented as log(D) for ProUbi10:CPK3-mEOS3.2, ProUbi10:CPK3-mEOS3.2 + PlAMV-GFP and ProUbi10:CPK3^CA^-mEOS3.2. Data were acquired from at least 6144 trajectories obtained in at least 15 cells over the course of three independent experiments. D. Box plots of the fraction of immobile proteins (log(D)<-2). Significant differences were revealed using a One-Way ANOVA followed by a Tukey’s multiple comparison test. Letters are used to discriminate between statistically different conditions (p<0.005). E. Mean square displacement (MSD) over time of fast (circle) or slow (triangle) diffusing ProUbi10:CPK3-mEOS3.2, ProUbi10:CPK3-mEOS3.2 + PlAMV-GFP, ProUbi10:CPK3^CA^-mEOS3.2. Standard error is displayed from three independent experiments. F. Voronoï tessellation illustration of ProUbi10:CPK3-mEOS3.2, ProUbi10:CPK3-mEOS3.2 + PlAMV-GFP and ProUbi10:CPK3^CA^-mEOS3.2. ND are circled in red. Scale bar = 2µm G. Distribution of the ND diameter of ProUbi10:CPK3-mEOS3.2, ProUbi10:CPK3-mEOS3.2 + PlAMV-GFP and ProUbi10:CPK3^CA^-mEOS3.2 H. Box plot representing the mean peak value of ND diameter extracted from the Gaussian fit of Figure 3G. No significant differences were revealed using a Kruskal-Wallis followed by a Dunn’s multiple comparison test. I. Boxplot of the proportion of detections found in ND of ProUbi10:CPK3-mEOS3.2, ProUbi10:CPK3-mEOS3.2 + PlAMV-GFP and ProUbi10:CPK3^CA^-mEOS3.2. Significant differences were revealed using a Kruskal-Wallis followed by a Dunn’s multiple comparison test. Letters are used to discriminate between statistically different conditions (p<0.005). J. Left: Confocal images of the surface view of *N. benthamiana* epidermal cells transiently expressing ProUbi10:CPK3^CA^-mRFP1.2 and infiltrated with either DMSO or 50 µg/mL fenpropimorph. Scale bar = 5µm; Right: Box plot of the mean Spatial Clustering Index (SCI) of CPK3^CA^. At least three experiments were performed, with at least 10 cells per experiment; statistical significance was determined using a Student *t*-test, ****: p < 0.0001. K. Left: Confocal images of the surface view of *N. benthamiana* epidermal cells transiently co-expressing ProUbi10:CPK3^CA^-mRFP1.2 with active or dead SAC1, mutated for its phosphatase activity. Scale bar = 5µm; Right: Box plot of the mean SCI of CPK3^CA^. At least three experiments were performed, with at least 10 cells per experiment; statistical significance was determined using a Student *t*-test, ****: p < 0.0001.

We analyzed the dynamics of ProUbi10:CPK3-mEOS3.2 and ProUbi10:CPK3^CA^-mEOS3.2 by spt-PALM (Figure 3B). We observed that the fraction of immobile ProUbi10:CPK3^CA^-mEOS3.2 molecules (log(D) < −2) is more abundant than for ProUbi10:CPK3-mEOS3.2 in absence of viral infection while no significant difference could be deciphered upon PlAMV infection (Figure 3C and D). In addition, MSD analysis showed that the motion of CPK3-mEOS3.2 mobile fraction is less confined than the one of CPK3^CA^-mEOS3.2 and CPK3-mEOS3.2 upon PlAMV infection (Figure 3E). Overall, this indicates that the mobile and immobile fraction of CPK3 is affected upon PlAMV infection and upon truncation of its auto-inhibitory domain, respectively.

Cluster analysis of CPK3 was performed using tessellation on the localization data obtained with spt-PALM (Figure 3F). Although we did not observe any significant differences in distribution of cluster sizes between all compared conditions(Figure 3G and H), CPK3^CA^ displayed a significantly higher proportion of proteins localized in ND compared to CPK3 (Figure 3I).

While the mechanism(s) governing the clustering of membrane proteins are not fully described, it is widely accepted that lateral organization involves – to some extent – protein-lipid interactions and lipid-lipid organization^10,11,38^. We observed that the integrity of ProUbi10:CPK3^CA^-mRFP1.2 organization relied on sterols and phosphoinositides. Indeed, treatment with fenpropimorph, a well-described inhibitor of sterol biosynthesis^39^, abolished CPK3^CA^ ND organization (Figure 3J), and the co-expression of ProUbi10:CPK3^CA^-mRFP1.2 with the yeast phosphatidylinositol-4-phosphate (PI4P)-specific phosphatase SAC1 targeted to the PM^40^, led to sparser and bigger domains (Figure 3K), which suggested that PI4P is not required for CPK3^CA^ ND formation but for its regulation.

All together these observations show that either removal of the auto-inhibitory domain or infection with PlAMV-GFP modifies CPK3 dynamics within the PM ; a mechanism which could be at stake for kinase activation.

### PlAMV infection induces an increase in REM1.2 PM diffusion

Group 1 REMs are one of the targets of CPK3 and we previously demonstrated that the restriction of PVX propagation by CPK3 overexpression depended on endogenous group 1 NbREMs^15^. Four REM isoforms belong to the group 1 in *Arabidopsis*: REM1.1, REM1.2, REM1.3 and REM1.4^41^. *REM1.2* and *REM1.3* are amongst the 10% most abundant transcripts in Arabidopsis leaves while REM1.1 was not detected in recently published leaf transcriptomes and proteomes^42^. Therefore, we focused on the three isoforms REM1.2, REM1.3 and REM1.4. REMs are described as scaffold proteins^43^, for which physiological function depends on the proteins they interact with and their phosphorylation status^44^. As recently described in^45^, REM1.2 and REM1.3 share 95% of their interactome, suggesting that they are functionally redundant. To address this, we isolated single T-DNA mutants *rem1.2*, *rem1.3* and *rem1.4* (SALK_117637.50.50.x, SALK_117448.53.95.x and SALK_073841.47.35, respectively) and crossed them to obtain the double mutant *rem1.2 rem1.3* and the triple mutant *rem1.2 rem1.3 rem1.4* (Figure 4 – figure supplement 1). We did not notice any obvious defects in the growth and development of seedlings and adult plants for the single, double and triple mutants, when grown under our conditions (Figure 4 – figure supplement 2). No difference in PlAMV-GFP propagation could be observed in the single mutants, compared to Col-0 (Figure 4A). However, the double mutant *rem1.2 rem1.3* showed a significant increase of infection foci area compared to Col-0, which was further enhanced in *rem1.2 rem1.3 rem1.4* triple KO mutant. Such additive effect of multiple mutations shows that REM1.2, REM1.3 and REM1.4 are functionally redundant regarding PlAMV cell-to-cell propagation. Finally, PlAMV-GFP systemic propagation was followed in whole plants every 3-4 days from 10 to 17 dpi and *rem1.2 rem1.3 rem1.4* displayed an increased infection surface of systemic leaves compared to Col-0 (Figure 4B and C), suggesting that group 1 REMs are involved in both local and systemic propagation of PlAMV-GFP.

**Figure 4.**
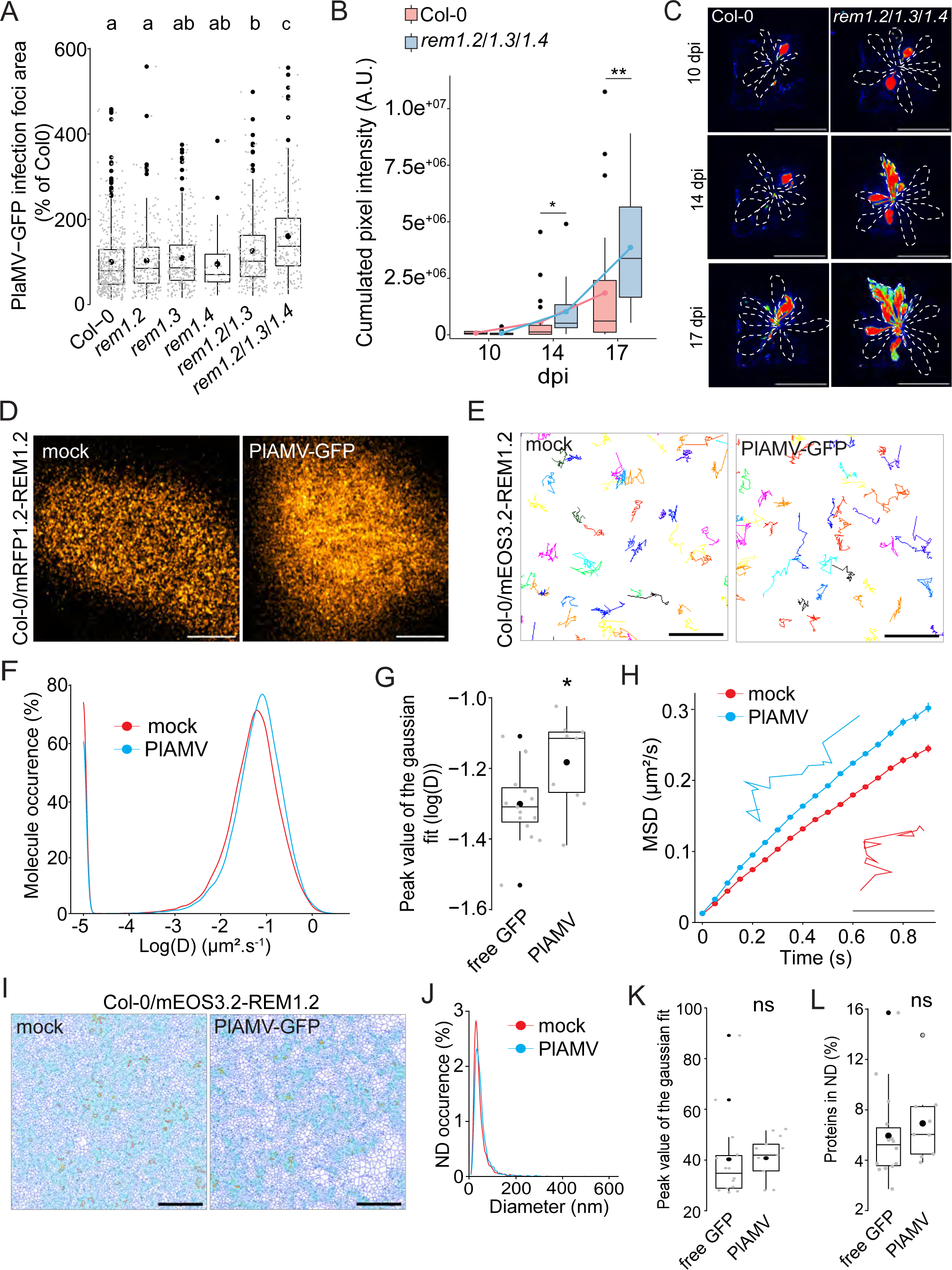
Group 1 REM hampers PlAMV-GFP cell-to-cell propagation and REM1.2 diffusion increases upon infection. A. Box plots of the mean area of PlAMV-GFP infection foci in *rem1.2, rem1.3, rem1.*4 single mutants along with *rem1.2 rem1.3* double mutant and *rem1.2 rem1.3 rem1.4* triple mutant. Three independent biological repeats were performed, with at least 36 foci per experiment and per genotype. Significant differences were revealed using a One-Way ANOVA followed by a Tukey’s multiple comparison test. Letters are used to discriminate between statistically different conditions (p<0.05). B. Box plots of the mean cumulated intensity measured in infected leaves in Col-0 and *rem1.2 rem1.3 rem1.4* during systemic viral propagation. Two independent experiments were conducted. Statistical significance of the difference between Col-0 and *rem1.2 rem1.3 rem1.4* at each time point was assessed using a Mann-Whitney test. *: p<0.05, **: p<0.01 C. Representative images of *A. thaliana* plants infected with PlAMV-GFP and imaged with a CCD Camera from 10 to 17 dpi. Systemic leaves are circled with a white dotted line. Multicolored scale is used to enhance contrast and ranges from blue (low intensity) to red (high intensity). Scale bar = 4 cm. D. Confocal images of the surface view of *A. thaliana* epidermal cells of the Col-0/ProUbi10:mRFP1.2-REM1.2, infiltrated either with free GFP (“mock”) or PlAMV-GFP. Scale bar = 5µm E. Representative trajectories of Col-0/ProUbi10:mEOS3.2-REM1.2 five days after infiltration with free GFP (“mock”) or PlAMV-GFP. Scale bar = 2µm F. Distribution of the diffusion coefficient (D), represented as log(D) for Col-0/ProUbi10:mEOS3.2-REM1.2 five days after infiltration with free GFP (“mock”) or PlAMV-GFP. Data were acquired from at least 28638 trajectories obtained from at least 16 cells over the course of three independent experiments. G. Box plot of the mean peak value extracted from the Gaussian fit of log(D) distribution. Significant difference was revealed using a Mann-Whitney test. *: p<0.05. H. Mean square displacement (MSD) over time of Col-0/ProUbi10:mEOS3.2-REM1.2 infiltrated either with free GFP (“mock”) or PlAMV-GFP. Representative trajectories extracted from Figure 4E illustrate each curve. Scale bar=1µm. I. Voronoi tessellation illustration of Col-0/ProUbi10:mEOS3.2-REM1.2 five days after infiltration with free GFP (“mock”) or PlAMV-GFP. ND are circled in red. Scale bar = 2µm J. Distribution of the ND diameter of Col-0/ProUbi10:mEOS3.2-REM1.2 five days after infiltration with free GFP (“mock”) or PlAMV-GFP. K. Box plot representing the mean peak value of ND diameter extracted from the Gaussian fit of Figure 4J. No significant difference was revealed using a Mann-Whitney test. L. Boxplot of the proportion of Col-0/ProUbi10:mEOS3.2-REM1.2 detections found in ND five days after infiltration with free GFP (“mock”) or PlAMV-GFP. No significant difference was revealed using a Mann-Whitney test.

Given their role in cell-to-cell viral movement, we checked whether group 1 REM expression was modified upon infection. RT-qPCR and western blots showed that neither transcripts nor protein levels were modified upon PlAMV infection (Figure 4 – figure supplement 3). Since REM1.2 and REM1.3 share a large part of their interactome and show functional redundancy regarding PlAMV infection, we decided to focus on REM1.2 for further investigations. Confocal imaging of the surface of epidermal cells of Col-0/ProUbi10:mRFP1.2-REM1.2 showed a rather heterogeneous distribution at the PM, although less striking than previously described when observed in root^31^ (Figure 4D). REM1.2 was next fused to mEOS3.2 and stably expressed in Col-0 to conduct spt-PALM (Figure 4E). The diffusion coefficient of ProUbi10:mEOS3.2-REM1.2 was significantly increased upon PlAMV infection (Figure 4F and G). Moreover, its MSD was increased to a similar extent as what was previously observed with StREM1.3 during a PVX infection^15^ (Figure 4H), although REM1.2 is overall more mobile than StREM1.3. Tessellation analysis of protein localization did not show any difference in ND organization, whether in size or regarding the enrichment of proteins in ND (Figure 4I-L).

Taken together, these results show that group 1 REMs are functionally redundant regarding their ability to hamper PlAMV propagation. PlAMV infection promoted an increased diffusion of REM1.2, in the same way as PVX did with StREM1.3. The conservation of such mechanism between plants of different families is indicative of its physiological importance.

### PlAMV-induced changes in REM1.2 and CPK3 plasma membrane dynamics are interdependent

Group 1 REMs from *A. thaliana* were previously identified as *in vitro* substrates of CPK3^19^. Moreover, untargeted immunoprecipitation experiments coupled to mass spectrometry identified CPK3 as an interactor of REM1.2 in *A. thaliana*^45^. We wanted to assess the functional link between CPK3 and group 1 REM in potexvirus propagation by knocking out CPK3 into the *rem1.2 rem1.3 rem1.4* mutant background. We isolated two independent CRISPR-generated *rem1.2 rem1.3 rem1.4 cpk3* #1 and *rem1.2 rem1.3 rem1.4 cpk3* #2 quadruple mutants (Figure 5 – figure supplement 1). We did not observe any developmental defect in these lines when grown under controlled conditions (Figure 5 – figure supplement 2). The analysis of PlAMV-GFP propagation showed that no significant additive effect could be observed between the quadruple mutant lines, *cpk3-2* and *rem1.2 rem1.3 rem1.4* (Figure 5A and 5B). This indicates that group 1 REMs and CPK3 function in the same signaling pathway to inhibit PlAMV propagation.

**Figure 5.**
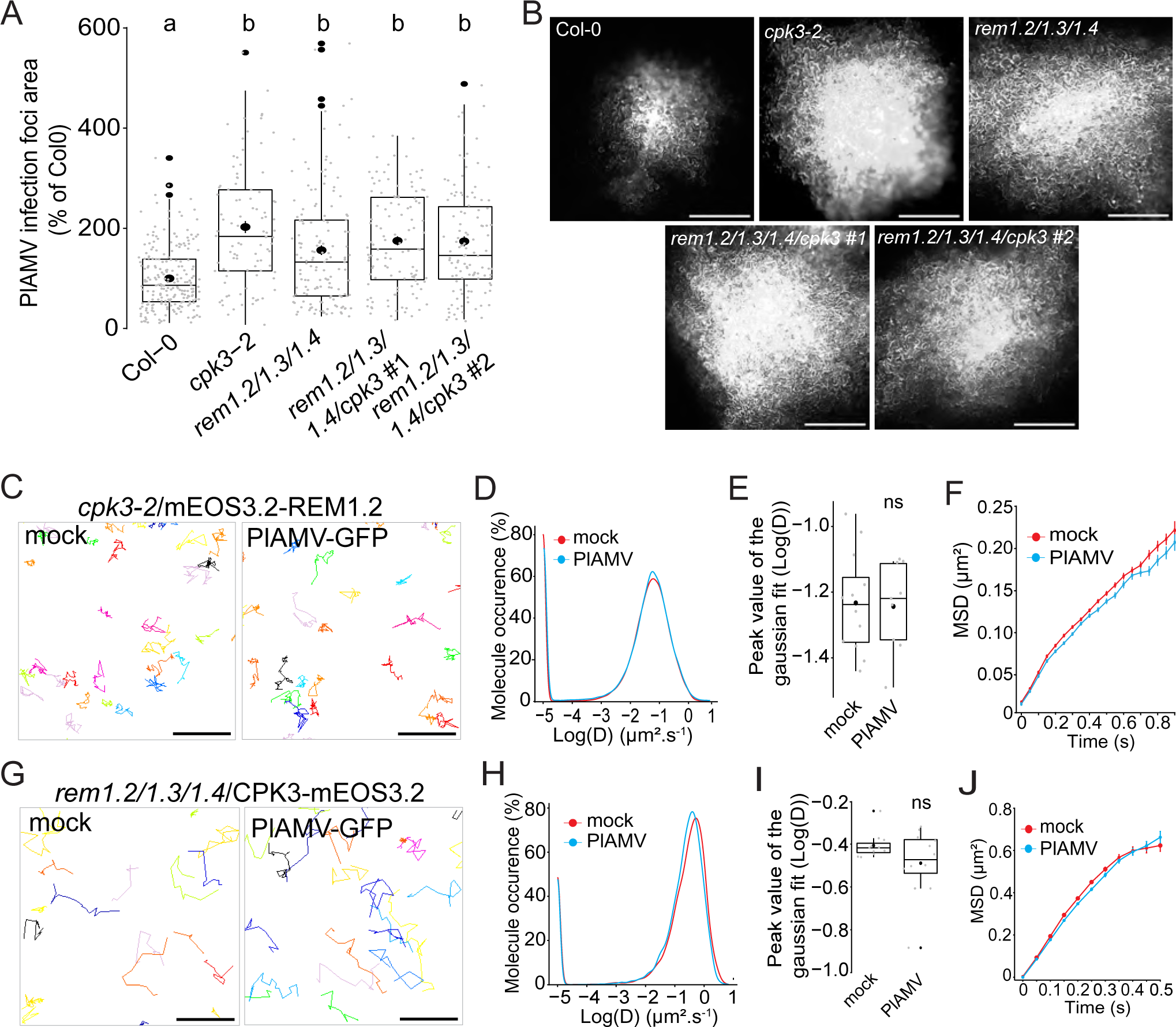
Group 1 REMs and CPK3 are in the same functional pathway and regulate each other PM diffusion upon PlAMV infection. A. Box plots of the mean area of PlAMV-GFP infection foci in *cpk3-2*, *rem1.2 rem1.3 rem1.4* triple mutant and *rem1.2 rem1.3 rem1.4/cpk3 #1 and #2* quadruple mutants. Three independent biological repeats were performed, with at least 23 foci per experiment and per genotype. Significant differences were revealed using a One-Way ANOVA followed by a Tukey’s multiple comparison test. Letters are used to discriminate between statistically different conditions (p<0.0001). B. Representative images of PlAMV-GFP infection foci at 5 dpi in the different mutant backgrounds. Scale bar = 500µm. C. Representative trajectories of *cpk3-2*/ProUbi10:mEOS3.2-REM1.2 five days after infiltration with either free GFP (“mock”) or PlAMV-GFP. Scale bar = 2µm. D. Distribution of the diffusion coefficient (D), represented as log(D) for *cpk3-2*/ProUbi10:mEOS3.2-REM1.2 five days after infiltration with either free GFP (“mock”) or PlAMV-GFP. Data were acquired from at least 20462 trajectories obtained from at least 11 cells over the course of three independent experiments. E. Box plot representing the mean peak value extracted from the Gaussian fit of the distribution of the diffusion coefficient (D) represented in Figure 5D. No significant difference was revealed using a Mann-Whitney test. F. Mean square displacement (MSD) over time of *cpk3-2*/ProUbi10:mEOS3.2-REM1.2 five days after infiltration with free GFP (“mock”) or PlAMV-GFP. G. Representative trajectories of *rem1.2 rem1.3 rem1.4*/ProUbi10:CPK3-mEOS3.2 five days after infiltration with free GFP (“mock”) or PlAMV-GFP. Scale bar = 2µm. H. Distribution of the diffusion coefficient (D), represented as log(D) for *rem1.2 rem1.3 rem1.4*/ProUbi10:CPK3-mEOS3.2 five days after infiltration with free GFP (“mock”) or PlAMV-GFP. Data were acquired from at least 11724 trajectories obtained from at least 10 cells over the course of three independent experiments. I. Box plot representing the mean peak value extracted from the Gaussian fit of the distribution of the diffusion coefficient (D) represented in Figure 5H. No significant difference was revealed using a Mann-Whitney test. J. Mean square displacement (MSD) over time of *rem1.2 rem1.3 rem1.4*/ProUbi10:CPK3-mEOS3.2 five days after infiltration with free GFP (“mock”) or PlAMV-GFP.

We wanted to know whether the increased diffusion of REM1.2 observed on PlAMV infection was dependent on CPK3. Using spt-PALM, we obtained the diffusion parameters of ProUbi10:REM1.2-mEOS3.2 expressed in the *cpk3-2* mutant background. Strikingly, we observed that both the diffusion coefficient and MSD of REM1.2 were not anymore affected during a viral infection (Figure 5C-F), showing that REM1.2 PM lateral diffusion upon PlAMV infection depends on CPK3. In a similar manner as in Col-0, REM1.2 clustering upon PlAMV infection in *cpk3-2* background did not display any difference to the mock-infected plants (Figure 5 – figure supplement 3).

Moreover, we wondered whether the reciprocal effect was true for the diffusion of CPK3 in the absence of group 1 REMs. Similarly, the diffusion coefficient and the MSD of mEOS3.2-CPK3 in *rem1.2 rem1.3 rem1.4* triple KO background remained the same during an infection compared to control condition (Figure 5G-J), unlike what was observed in a Col-0 background (Figure 2D-G). This result indicated that the confinement of CPK3 proteins upon viral infection depended on the presence of group 1 REMs. Moreover, contrarily to the Col-0 background, the *rem1.2 rem1.3 rem1.4* displayed reduced protein concentration upon PlAMV infection (Figure 5 – figure supplement 3). This showed that group 1 REMs might play a role in CPK3 domain regulation upon viral infection.

As CPK3^CA^ was shown to inhibit potexvirus propagation more efficiently than the full-length protein^15^, we tested CPK3^CA^ ability to alter REM diffusion (Figure 5 – figure supplement 4). Upon transient co-expression with CPK3^CA^, REM diffusion was significantly increased. As shown in Figure 2B, CPK membrane anchor is crucial for its role in viral propagation. We tested if CPK3^CA-G2A^ was still able to hinder REM diffusion, which was not the case. All these data support a specific role of PM-bound CPK3 in REM increased diffusion upon viral infection.

We then investigated whether CPK3 and REM would colocalize in absence or presence of the virus. Using confocal microscopy, we showed that they randomly colocalized in both situations (Figure 5 – figure supplement 5), the interaction between the kinase and its substrate probably occurring in a narrow spatiotemporal window.

Taken all together, those results show a strong inter-dependence of group 1 REMs and CPK3 both in their physiological function and in their PM lateral diffusion upon PlAMV infection.

## DISCUSSION

### CPK3 specific role in viral immunity is supported by its PM organization

Although calcium-mediated signaling is suspected to be involved in viral immunity, only few calcium-modulated proteins are described as playing a role in viral propagation^15,46^. We showed here the crucial role of CPK3 over other immunity-related CPK isoforms^24,25^ by a reverse genetic approach in *Arabidopsis*. The precise role of CPK3 in viral immunity remains to be determined. CPK3 phosphorylates actin depolymerization factors to modulate the actin cytoskeleton^21,47^, a key player in host-pathogen interaction^48^. In particular, potexviruses induce the remodeling of the actin cytoskeleton to organize the key steps of their cycle, whether it is replication, intra-cellular movement or cell-to-cell propagation^49,50^. PlAMV replication and/or movement could be affected by CPK3-mediated alteration of the cytoskeleton mesh.

The specificity of a calcium-dependent kinase in a given biological process is determined by its expression pattern, subcellular localization and substrate specificity^27^ Controlled subcellular localization ensures proximity with either stimulus or substrate and here we showed that, similarly to *Arabidopsis* CPK6^52^ and *Solanum tuberosum* CPK5^53^, the disruption of CPK3 membrane anchor led to a loss-of-function phenotype (Figure 2B). Interestingly, we observed a reduction in CPK3 PM diffusion upon PlAMV infection, suggesting that not only membrane localization but also protein organization at the PM is important during viral immunity. Moreover, PlAMV-induced CPK3 PM confinement was reminiscent of the diffusion parameters displayed by CPK3^CA^, hinting that viral infection, kinase activation and lateral diffusion are linked. However, it is necessary to remain careful as a truncated protein deprived of its auto-inhibitory domain does not reflect the controlled and context-dependent activation of CPK3. Indeed, stable expression of CPK3^CA^ was previously shown to be lethal^37^. CPK3^CA^ ever-activated state might lead to stable or unspecific interaction with protein partners along with erratic phosphorylation of substrates, which could explain the PM domains formed by CPK3^CA^ at the PM.

CPK3 nanoscale dynamics upon viral infection might offer another layer of specificity to convey the appropriate response to a given stimulus by ensuring proximity with specific regulators or substrates. It would be interesting to explore whether the reduction of CPK3 diffusion observed upon PlAMV infection is specific to this virus or if it can be extended to other viral species, genera and even pathogens. Indeed, it was recently shown that CPK3 transcription was enhanced upon infection by viruses from different genera^54^. Moreover, CPK3 regulates herbivore responses by phosphorylating transcription factors^20^, is activated by flg22 in protoplasts^19^ and is proposed to be the target of a bacterial effector to disrupt immune response^21^. Finally, the diffusion and clustering of other PM-localized CPKs could be investigated as no experimental data exist yet regarding their PM nano-organization. It would be especially relevant to describe these parameters for the CPK isoforms phosphorylating ND-organized NADPH oxidases^14,25,28^.

### Potential functions for REM1.2 increased diffusion upon PlAMV

REM proteins display a wide range of physiological functions and are proposed to function as scaffold proteins^43,44,55^. Group 1 REMs have been shown to be involved in viral immunity, playing apparent contradictory roles depending on the virus genera^8,44,56^. Herein, we show through a combination of genetic and biochemical analysis that the three most expressed isoforms of group 1 *Arabidopsis* REMs were functionally redundant in inhibiting PlAMV propagation (Figure 4A). REMs are anchored at the PM through their C-terminal sequence^44,56–58^ but despite displaying a similar PM attachment, potato StREM1.3 is static and forms well defined membrane compartments^57^ while *Arabidopsis* REM1.2 appeared mobile in leaves, with small and potentially short-lived membrane domains of around 70nm (Figure 4F-J). Beyond clear organizational differences, both REM1.2 and StREM1.3 showed an increased mobility upon viral infection in *N. benthamiana* and *Arabidopsis*^15^ (Figure 4E-H), which is at contrast with the canonical model linking protein activation with its stabilization into ND^11,59^. Particularly, REM1.2 was recently shown to form ND upon elicitation by a bacterial effector^13^ or upon exposition to bacterial membrane structures^60^. Conservation across plant and virus species of such increased PM diffusion of group 1 REMs indicates that this specific mechanism is crucial for plant response to potexviruses, although its role remains to be deciphered. It was suggested, in the context of tobacco rattle virus infection, that REM1.2 clusters led to an increased lipid order of the PM and a morphological modification of plasmodesmata (PD), inducing a decrease of PD permeability^31^. As REM1.2 does not accumulate at PD upon PlAMV infection (Figure 4 – figure supplement 4), it might indirectly influence PD permeability through mechanisms yet to be discovered. We previously showed that StREM1.3 modulated the formation of lipid phases *in vitro*^61^ while *Medicago truncatula* SYMREM1 was recently shown to stabilize membrane topology changes in protoplasts^61^. Therefore, REM1.2 increased diffusion upon viral infection might alter lipid organization, with consequences on PM and PD-localized proteins^62^. Based on REM’s putative scaffolding role^43,63^, its increased PM diffusion might modify the stability of REM-supported complexes and induce subsequent signaling, similarly to the way *Oryza sativa* REM4.1 orchestrates the balance between abscisic acid and brassinosteroid pathways by interacting with different protein kinases^64^.

### A PM-localized mechanism involved in hampering viral propagation

Increasing evidence supports the role of PM proteins in plant antiviral mechanism^65^. However, while the nano-organization of PM proteins involved in bacterial or fungal immunity begins to be addressed^13,60,66,67^, there is still only scarce information in a viral infection context. Our observations pointed towards an interdependence of REM and CPK3 in their PM diffusion upon PlAMV infection. In particular, CPK3-dependent REM increased diffusion in a potexvirus infection is consistent with our previously reported PM diffusion increase of StREM1.3 phosphomimetic mutant, reminiscent of StREM1.3 behavior upon viral infection^15^. Moreover, we recently published that *in vitro* CPK3-phosphorylated StREM1.3 presented disrupted domains on model membranes compared to the non-phosphorylated protein^16^. The data presented in this paper further support the predominant role of CPK3 in viral-induced REM1.2 diffusion (Figure 5C-F). Since the actin cytoskeleton was shown to favor nanometric scale ND including those of group 1 REM^68^, CPK3 mediated regulation of the actin cytoskeleton could regulate REM1.2 PM nanoscale organization. We also discovered the essential role of group 1 REMs in the lateral organization of CPK3 upon viral infection (Figure 5G-J), which is further supported by the difference between CPK3 clustering parameters when expressed in *N. benthamiana* or in *Arabidopsis*: CPK3 was enriched in ND upon PlAMV infection when transiently expressed in *N. benthamiana* but not in *Arabidopsis* (Figure 2H, 2K, 3F and 3I). The discrepancy between both species could be linked to *N. benthamiana* REM1 ability to form stable ND, discernable by confocal microscopy^5^ while we observed small and unstable domains of REM1.2 in *Arabidopsis* leaves (Figure 4I-K). Moreover, CPK3 ND organization was disrupted in *rem1.2 rem1.3 rem1.4* triple KO background upon PlAMV infection (Figure 5 – figure supplement 3H), indicating that group1 REMs are crucial for the spatial organization of CPK3 during a viral infection. Furthermore, the sensitivity of CPK3^CA^ domains to the same lipid inhibitors as StREM1.3^57^ (Figure 3J,K) hints that REM1.2 lipid environment might be required for CPK3 nanoscale organization.

Although the data obtained here do not allow us to determine the sequence of events orchestrating REM1.2 and CPK3 dynamic interaction, hypotheses can be formulated (Figure 6). REM and CPK3 interact in the absence of any stimulus^15,45^ but as they display drastically different diffusion parameters (Figure 2E, Figure 4F), likely only a small part of proteins interact at basal state. PlAMV infection induces a REM-dependent confinement of CPK3 (Figure 2 and Figure 5). Whether CPK3 confinement is concomitant to its activation remains to be confirmed. Such modification of CPK3 mobility might result in an increase probability of interaction with REM1.2, which might lead to increased phosphorylation events and a subsequent increase in mobility of REM1.2. In addition to the above-mentioned putative roles of REM’s increased diffusion, such “kiss-and-go” mechanism between a kinase and its substrate might also be considered as a negative feedback loop to hamper constitutive activation of the system. It was recently shown that the PM organization of a receptor-like kinase and its co-receptor relied on the expression of a scaffold protein^67,69^, REM might go beyond the role of substrate and be essential to maintain the required PM environment of its cognate kinase to ensure proper signal transduction. Although the complexity of potexvirus’ lifecycle as well as technical limitations impair a thorough challenge of this hypothesis, our data support the significance of fine PM compartmentalization and spatio-temporal dynamics in signal transduction upon viral infection in plants while it explores new concepts underlying plant kinases PM organization.

**Figure 6.** Proposed “kiss-and-run” model describing REM1.2 and CPK3 interdependence. In healthy conditions, REM1.2 displays a smaller mobility at the PM than CPK3. Random interactions between the two proteins can occur. Upon PlAMV infection, CPK3 shows a REM-dependent confined behavior which might be concomitant to its activation. Simultaneously, a PlAMV infection leads to a CPK3-dependent increased diffusion of REM1.2, which might be a consequence of its phosphorylation by CPK3.

## ACKNOWLEDGEMENTS

We thank Thierry Mauduit and Christophe Higelin (HPE Greenhouse, INRAe) for plant culture. We thank the Bordeaux Imaging Center, part of the National Infrastructure France-BioImaging supported by the French National Research Agency (ANR-10-INBS-04). This work was supported by the French National Research Agency (grant no. ANR-19-CE13-0021 to SGR, SM, VG, MB) and the German Research Foundation (DFG) grant CRC1101-A09 to JG, the IPS2 benefits from the support of the LabEx Saclay Plant Sciences-SPS (ANR-10-LABX-0040-SPS). This study received financial support from the French government in the framework of the IdEX Bordeaux University “Investments for the Future” program / GPR “Bordeaux Plant Sciences”.

## AUTHORS CONTRIBUTION

Conceptualization: M.-D.J., S.M, V.G.

Investigation: M-D. J, A.-F. D., M.B., N.B.A., M.R., T.R., V. W.-B., J.H., D.L., G.S., J.A

Resources: M.-D.J., M.B., V.W.-B., J.H., D.L., N.B.A., Y.-J.L., B.D., V.C., N.G., J.-L.G., Y.Y. T.O.

Writing (Original Draft): M.-D.J., S.M.

Writing (Review and Editing): M.-D.J., M.B., N.A., V.W.-B., J.A., V.C., J.-L.G., S.G.-R., J.G., S.M, V.G.

Visualization: M.D.J, A-F.D.

Funding acquisition: S.M., V.G., S.G.-R., T.O.

## MATERIAL AND METHODS

### Plant culture

*Nicotiana benthamiana* plants were cultivated in controlled conditions (16 h photoperiod, 25 °C). Proteins were transiently expressed via *Agrobacterium tumefaciens* as previously described^57^. The agrobacteria GV3101 strain was cultured at 28°C on appropriate selective medium depending on constructs carried. Plants were observed between 2 and 5 days after infiltration depending on experiments.

Sterilized *Arabidopsis. thaliana* seeds were germinated on ½ MS plates supplemented with 1% sucrose. 10-day-old seedlings were transferred to soil and grown under short day conditions (8 h light/ 16 h dark).

### Cloning

REM1.2, CPK3 and CPK3^CA^ sequences were previously published^15^. CPK3^K107M^ was generated by site-directed mutagenesis using CPK3 as a template and primers specified in the Supplemental Table 1.

All vectors built for this project, except for CRISPR and some protein production, were generated using multisite Gateway cloning strategies (www.lifetechnologies.com) with pDONR P4-P1r, pDONR P2R-P3, pDONR221 as entry vectors. pLOK180_pR7m34g^71^ was used as a destination vector for plant expression. pGEX-2T (GE Healthcare, N-terminal fusion with GST) was used for CPK protein production in bacteria for previously reported constructs^72^ (CPK2/3/5/11). CPK1, CPK3^K107M^ and CPK6 were cloned into pGEX-3X-GW using the gateway cloning system and pDONR207 as entry vector. The N-terminal 118 amino acids of REM1.2 (REM1.2^1–118^) was synthetized with optimized codons for bacterial expression by GenScript (genscript.com) and cloned into pET24a (C-terminal fusion with a 6-histidine tag) between Nde1 and XhoI.

To generate CRISPR lines, sgRNAs targeting the N-terminus of *CPK3* gene were selected using CRISPR-P 2.0 website^73^ (http://crispr.hzau.edu.cn/CRISPR2) and cloned into pHEE401 backbone^74^ carrying the gene coding for the zCas9 enzyme under the control of Egg cell promoter using the Golden Gate cloning method.

All constructs were propagated using the NEB10 *E. coli* strain (New England Biolabs). Primers used for cloning are detailed in Supplemental Table 1.

### Plant lines generation

T-DNA insertion mutants *rem1.2* (salk_117637.50.50.x), *rem1.3* (salk_117448.53.95.x) and *rem1.4* (SALK_073841.47.35) were provided by the ABRC. *rem1.2 rem1.3* double mutant was generated by crossing the respective T-DNA inserted parental plants, *rem2/rem1.3/rem1.4* was created by crossing *rem1.2 rem1.3* with *rem1.4.* The *cpk5 cpk6* (sail_657_C06, salk_025460) and *cpk5 cpk6 cpk11* (sail_657_C06, salk_025460, salk_054495) were described previously^24^. The quadruple mutant *cpk3 cpk5 cpk6 cpk11* was described previously^26^. *cpk1 cpk2* (salk_096452, salk_059237) and *cpk1 cpk2 cpk5 cpk6* (salk_096452, salk_059237, sail_657_C06, salk_025460) were described previously^25^. CPK3 T-DNA insertion lines *cpk3-1* (salk_107620) and *cpk3-2* (salk_022862) were obtained from Julian Schroeder^32^ and Bernhard Wurzinger^19^, respectively. All mutants are in the same genetic ecotype Columbia Col-0. All plants were genotyped using primers indicated in the Supplemental Table 1.

Pro35S:CPK3-HA #16.2, *cpk3-2*/Pro35S:CPK3-myc and *cpk3-2*/Pro35S:CPK3^K107M^-myc used for viral propagation were previously published^21,72^. Pro35S:CPK3-HA #8.2 was generated at the same time as Pro35S:CPK3-HA #16.2, and protein expression was confirmed (Figure 1 – figure supplement 3).

Col-0 plants were floral dipped with either ProUbi10:CPK3-mRFP1.2, ProUbi10:CPK3-mEOS3.2 or ProUbi10:mEOS3.2-REM1.2. *cpk3-2* plants were floral dipped with either ProUbi10:CPK3-mRFP1.2, ProUbi10:CPK3^G2A^-mRFP1.2 or ProUbi10:mEOS3.2-REM1.2. *rem1.2 rem1.3 rem1.4* plants were floral dipped with ProUbi10:CPK3-mEOS3.2. Col-0/ProREM1.2:YFP-REM1.2^75^ was transformed with ProUbi10:CPK3-mTagBFP2. Transformed seeds were selected based on the seedcoat RFP fluorescence.

For CRISPR-mediated site mutagenesis of CPK3, *rem1.2 rem1.3 rem1.4* was transformed by floral dip with the plasmid carrying z*Cas9* encoding gene and the sgRNA. Transformed candidates were selected on hygromycin and grown for seed collection. Harvested seeds were grown and a leaf sample was harvested for genomic DNA extraction. PCR were performed to amplify the targeted region, and CRISPR-induced mutations were screened using capillary electrophoresis. Mutated candidates were sent to sequencing to obtain homozygous lines for the mutation and then backcrossed with *rem1.2 rem1.3 rem1.4* to remove the Cas9. Primers used for CRISPR screening are listed in Supplemental Table 1.

### Local viral propagation assay

Viral propagation assays were performed using PlAMV-GFP, an agroinfiltrable GFP-tagged infectious clone of PlAMV ^22^*. Agrobacterium tumefaciens* strain GV3101 carrying PlAMV-GFP was infiltrated on 3-week-old *A. thaliana* plants at OD_600nm_ = 0.2. Viral spreading was tracked using Axiozoom V16 macroscope system 5 days after infection. Infection foci were automatically analyzed using the Fiji software (http://www.fiji.sc/) via a homemade macro. The statistical significance was assessed using a two-way ANOVA, followed by a Tukey’s multiple comparison test.

### Systemic viral propagation assay

At 3-week-old, leaves were infiltrated with *Agrobacterium tumefaciens* GV3101 strain carrying PlAMV-GFP vector. Infection was followed every three-four days from the 10^th^ day of infection to the 17^th^ using a closed GFP-CAM FX 800-0/1010 GFP camera and the Fluorcam7 software (Photon System Instruments, Czech Republic; https://fluorcams.psi.cz/). Image analysis was performed using Fiji software (https://fiji.sc/). Integrated density of the fluorescence of systemic leaves was measured. Two independent experiments with at least 30 plants per genotype were performed. Statistical significance was determined with a Mann-Whitney test or a Kruskal-Wallis followed by a Dunn’s multiple comparison test, depending on the number of conditions.

### Confocal microscopy

Live imaging was performed using a Zeiss LSM 880 confocal laser scanning microscopy system using either an oil-immersion PL-APO 40x or 68x (NA = 1.4) objective coupled with an AiryScan detector for the latter. mRFP1.2 fluorescence was observed using an excitation wavelength of 561 nm and an emission wavelength of 579 nm. Acquisition parameters remained the same across experiments for SCI quantification. The SCI was calculated as previously described ^57^. Briefly, 10 µm lines were plotted across the samples and the SCI was calculated by dividing the mean of the 5% highest values by the mean of 5% lowest values. Three lines were randomly plotted per cell. Three independent experiments were done, on at least 10 cells each time. Fenpropimorph (10µg/mL) or DMSO was infiltrated 24 hours before observation.

### spt-PALM microscopy

*N. benthamiana* and *A. thaliana* epidermal cells were imaged at RT. Samples of leaves of 3-week-old plants stably or transiently expressing mEOS3.2-tagged constructs were mounted between a glass slide and a cover slip in a drop of water to avoid dehydration. With the exception of ProUbi10:mEOS3.2-REM1.2 transiently expressed in *N. benthamiana*, image acquisitions were done on an inverted motorized microscope Nikon Ti Eclipse equipped with a 100Å∼ oil-immersion PL-APO objective (NA = 1.49), a TIRF arm and a sCMOS Camera FUsion BT (Hamamatsu). Laser angle was adjusted to obtain the highest signal-to-noise ratio while laser power of a 405nm and 561nm laser, respectively to activate and image mEOS3.2, was adjusted to obtain a sufficient concentration of individual particles. Particle localization, tracks reconstructions, diffusion coefficient and MSD parameters were obtained using PALMtracer, as previously described^57^. The diffusion coefficient was calculated from the four first points of the MSD. Three independent experiments were conducted for each tested condition. Tessellation analysis was conducted using SR-Tesseler as previously described^34,57^. ND were considered to be at least 32 nm², to contain at least 5 particles and to have a particle density twice the average particle density within the sample.

For ProUbi10:mEOS3.2-REM1.2 transiently expressed in *N. benthamiana* in the presence or absence of ProUbi10:CPK3^CA^-mVenus or ProUbi10:CPK3^CAG2A^-mVenus, images were acquired on a custom-built platform, as described previously^76^. Particle localization, tracks reconstructions, diffusion coefficient and MSD parameters were obtained using OneFlowTraX^76^. Particle localization was done using the following parameters: photon/ADU conversion of 0.48, an offset of 100, a pixel size of 100 nm, a filter size of 1.2, a cut-off of 3 and a PSF of 7. Track reconstruction was done with a maximum linking distance of 200 nm, a maximum gap frame of 4. Tracks smaller than 8 localizations were filtered out and the diffusion coefficient was calculated from the 2nd to 5th data points.

### RT-qPCR

3-week-old *A. thaliana* leaves were infiltrated with PlAMV-GFP at final OD_600nm_ = 0.2. Leaf samples were harvested 7 days after infiltration and immediately frozen. RNA extraction was done using Qiagen Plant Mini Kit and was followed by a DNase treatment. cDNA was produced from the extracted RNA using Superscript II enzyme from Invitrogen. RT-qPCR was performed on obtained cDNA using the iQ™ SYBR® Green supermix (BioRad) on the iQ iCycler thermocycler (BioRad). The transcript abundance in samples was determined using a comparative threshold cycle method and was normalized to actin expression. Statistical differences were determined using a Mann-Whitney test. Primers used for RT-qPCR are listed in Supplemental Table 1.

### Production of recombinant proteins and *in vitro* kinase assay

GST-CPK proteins were produced in BL21(DE3)pLys and purified as previously reported^72^. 6His-AtREM1.2^1–118^ was produced like GST-CPK and purified on Protino® Ni-TED column following manufacturer’s instructions (Macherey-Nagel). After elution with 40-250 mM imidazole, proteins were dialyzed overnight in 30 mM Tris HCl pH 7.5, 10% glycerol. Kinase assay was performed as previously described^15^ using 400 ng recombinant GST-CPK and 1-2 µg substrate (6His-AtREM1.2^1–118^ or histone).

### Production of CPK3 antibodies

Polyclonal antibodies against *Arabidopsis* CPK3 were raised in rabbit by Covalab (France) using purified recombinant GST-CPK3. The antibodies were purified from rabbit serum by affinity chromatography on CH-Sepharose 4B (GE Healthcare) coupled to 6His-CPK3. To produce the recombinant 6His-CPK3 and GST-CPK3 proteins, the Arabidopsis CPK3 cDNA was cloned into the expression vectors pDEST17 and pDEST15 (Invitrogen), respectively. Expression of 6His-CPK3 was induced in *Escherichia coli* strain BL21-AI (Invitrogen) with 0.2% (m/v) arabinose and the recombinant protein was affinity purified using Ni-NTA agarose (Qiagen). For GST-CPK3, protein expression in *E.coli* Rosetta cells (Novagen) was induced with 0.5 mM IPTG (isopropyl-β-D-thiogalactopyranoside) and recombinant GST-CPK3 was purified by Glutathione Sepharose 4 Fast Flow chromatography (GE Healthcare) as described by the manufacturer.

### Western Blots

Protein samples were extracted from *A. thaliana* leaf tissue using 2X Laemmli buffer or in a buffer containing 50 mM Tris HCl pH 7.5, 5 mM EDTA, 5 mM EGTA, 1X anti-protease cocktail [Roche], 1% Triton X-100, 2 mM DTT. Proteins were transferred to PVDF and detected with polyclonal antibodies raised against CPK3 or REM1.2/REM1.3^62^, followed by incubation with secondary anti-rabbit HRP-conjugated antibodies (Sigma).

## REFERENCES

1. Hanssen, I.M., and Thomma, B.P.H.J. (2010). Pepino mosaic virus: a successful pathogen that rapidly evolved from emerging to endemic in tomato crops. Mol. Plant Pathol. 11, 179–189. 10.1111/J.1364-3703.2009.00600.X.

2. Zorzatto, C., MacHado, J.P.B., Lopes, K.V.G., Nascimento, K.J.T., Pereira, W.A., Brustolini, O.J.B., Reis, P.A.B., Calil, I.P., Deguchi, M., Sachetto-Martins, G., et al. (2015). NIK1-mediated translation suppression functions as a plant antiviral immunity mechanism. Nature 520, 679. 10.1038/NATURE14171.

3. Ngou, B.P.M., Ding, P., and Jones, J.D.G. (2022). Thirty years of resistance: Zig-zag through the plant immune system. Plant Cell 34, 1447–1478. 10.1093/PLCELL/KOAC041.

4. Cheng, G., Yang, Z., Zhang, H., Zhang, J., and Xu, J. (2020). Remorin interacting with PCaP1 impairs Turnip mosaic virus intercellular movement but is antagonised by VPg. New Phytol. 225, 2122–2139. 10.1111/NPH.16285.

5. Fu, S., Xu, Y., Li, C., Li, Y., Wu, J., and Zhou, X. (2018). Rice Stripe Virus Interferes with S-acylation of Remorin and Induces Its Autophagic Degradation to Facilitate Virus Infection. Mol. Plant 11, 269–287. 10.1016/J.MOLP.2017.11.011.

6. Ma, T., Fu, S., Wang, K., Wang, Y., Wu, J., and Zhou, X. (2022). Palmitoylation Is Indispensable for Remorin to Restrict Tobacco Mosaic Virus Cell-to-Cell Movement in Nicotiana benthamiana. Viruses 14. 10.3390/V14061324.

7. Raffaele, S., Bayer, E., Lafarge, D., Cluzet, S., Retana, S.G., Boubekeur, T., Castel, N.L., Carde, J.P., Lherminier, J., Noirot, E., et al. (2009). Remorin, a solanaceae protein resident in membrane rafts and plasmodesmata, impairs potato virus X movement. Plant Cell 21, 1541–1555. 10.1105/tpc.108.064279.

8. Rocher, M., Simon, V., Jolivet, M.D., Sofer, L., Deroubaix, A.F., Germain, V., Mongrand, S., and German-Retana, S. (2022). StREM1.3 REMORIN Protein Plays an Agonistic Role in Potyvirus Cell-to-Cell Movement in N. benthamiana. Viruses 14. 10.3390/V14030574.

9. Son, S., Oh, C.J., and An, C.S. (2014). Arabidopsis thaliana remorins interact with SnRK1 and play a role in susceptibility to beet curly top virus and beet severe curly top virus. Plant Pathol. J. 30, 269–278. 10.5423/PPJ.OA.06.2014.0061.

10. Gronnier, J., Gerbeau-Pissot, P., Germain, V., Mongrand, S., and Simon-Plas, F. (2018). Divide and Rule: Plant Plasma Membrane Organization. Trends Plant Sci. 23, 899–917. 10.1016/j.tplants.2018.07.007.

11. Jaillais, Y., and Ott, T. (2020). The nanoscale organization of the plasma membrane and its importance in signaling: A proteolipid perspective. Plant Physiol. 182, 1682–1696. 10.1104/PP.19.01349.

12. Platre, M.P., Bayle, V., Armengot, L., Bareille, J., del Mar Marquès-Bueno, M., Creff, A., Maneta-Peyret, L., Fiche, J.B., Nollmann, M., Miège, C., et al. (2019). Developmental control of plant Rho GTPase nano-organization by the lipid phosphatidylserine. Science (80-. ). 364, 57–62. 10.1126/science.aav9959.

13. Ma, Z., Sun, Y., Zhu, X., Yang, L., Chen, X., and Miao, Y. (2022). Membrane nanodomains modulate formin condensation for actin remodeling in Arabidopsis innate immune responses. Plant Cell 34, 374–394. 10.1093/PLCELL/KOAB261.

14. Smokvarska, M., Francis, C., Platre, M.P., Fiche, J.B., Alcon, C., Dumont, X., Nacry, P., Bayle, V., Nollmann, M., Maurel, C., et al. (2020). A Plasma Membrane Nanodomain Ensures Signal Specificity during Osmotic Signaling in Plants. Curr. Biol. 30, 4654–4664.e4. 10.1016/j.cub.2020.09.013.

15. Perraki, A., Gronnier, J., Gouguet, P., Boudsocq, M., Deroubaix, A.-F., Simon, V., German-Retana, S., Legrand, A., Habenstein, B., Zipfel, C., et al. (2018). REM1.3’s phospho-status defines its plasma membrane nanodomain organization and activity in restricting PVX cell-to-cell movement. PLoS Pathog. 14. 10.1371/journal.ppat.1007378.

16. Legrand, A., G.-Cava, D., Jolivet, M.-D., Decossas, M., Lambert, O., Bayle, V., Jaillais, Y., Loquet, A., Germain, V., Boudsocq, M., et al. (2022). Structural determinants of REMORIN nanodomain formation in anionic membranes. Biophys. J. 10.1016/J.BPJ.2022.12.035.

17. Martinière, A., Fiche, J.B., Smokvarska, M., Mari, S., Alcon, C., Dumont, X., Hematy, K., Jaillais, Y., Nollmann, M., and Maurel, C. (2019). Osmotic stress activates two reactive oxygen species pathways with distinct effects on protein nanodomains and diffusion. Plant Physiol. 179, 1581–1593. 10.1104/pp.18.01065.

18. Gournas, C., Gkionis, S., Carquin, M., Twyffels, L., Tyteca, D., and André, B. (2018). Conformation-dependent partitioning of yeast nutrient transporters into starvation-protective membrane domains. Proc. Natl. Acad. Sci. U. S. A. 115, E3145–E3154. 10.1073/PNAS.1719462115/VIDEO-7.

19. Mehlmer, N., Wurzinger, B., Stael, S., Hofmann-Rodrigues, D., Csaszar, E., Pfister, B., Bayer, R., and Teige, M. (2010). The Ca2+-dependent protein kinase CPK3 is required for MAPK-independent salt-stress acclimation in Arabidopsis. Plant J. 63, 484–498. 10.1111/j.1365-313X.2010.04257.x.

20. Kanchiswamy, C.N., Takahashi, H., Quadro, S., Maffei, M.E., Bossi, S., Bertea, C., Zebelo, S.A., Muroi, A., Ishihama, N., Yoshioka, H., et al. (2010). Regulation of Arabidopsis defense responses against Spodoptera littoralis by CPK-mediated calcium signaling. BMC Plant Biol. 10, 97. 10.1186/1471-2229-10-97.

21. Lu, Y.J., Li, P., Shimono, M., Corrion, A., Higaki, T., He, S.Y., and Day, B. (2020). Arabidopsis calcium-dependent protein kinase 3 regulates actin cytoskeleton organization and immunity. Nat. Commun. 11. 10.1038/s41467-020-20007-4.

22. Minato, N., Komatsu, K., Okano, Y., Maejima, K., Ozeki, J., Senshu, H., Takahashi, S., Yamaji, Y., and Namba, S. (2014). Efficient foreign gene expression in planta using a plantago asiatica mosaic virus-based vector achieved by the strong RNA-silencing suppressor activity of TGBp1. Arch. Virol. 10.1007/s00705-013-1860-y.

23. Yamaji, Y., Maejima, K., Komatsu, K., Shiraishi, T., Okano, Y., Himeno, M., Sugawara, K., Neriya, Y., Minato, N., Miura, C., et al. (2012). Lectin-mediated resistance impairs plant virus infection at the cellular level. Plant Cell 24, 778–793. 10.1105/tpc.111.093658.

24. Boudsocq, M., Willmann, M.R., McCormack, M., Lee, H., Shan, L., He, P., Bush, J., Cheng, S.H., and Sheen, J. (2010). Differential innate immune signalling via Ca 2+ sensor protein kinases. Nature 464, 418–422. 10.1038/nature08794.

25. Gao, X., Chen, X., Lin, W., Chen, S., Lu, D., Niu, Y., Li, L., Cheng, C., McCormack, M., Sheen, J., et al. (2013). Bifurcation of Arabidopsis NLR Immune Signaling via Ca2+-Dependent Protein Kinases. PLOS Pathog. 9, e1003127. 10.1371/JOURNAL.PPAT.1003127.

26. Guzel Deger, A., Scherzer, S., Nuhkat, M., Kedzierska, J., Kollist, H., Brosché, M., Unyayar, S., Boudsocq, M., Hedrich, R., and Roelfsema, M.R.G. (2015). Guard cell SLAC1-type anion channels mediate flagellin-induced stomatal closure. New Phytol. 10.1111/nph.13435.

27. Yip Delormel, T., and Boudsocq, M. (2019). Properties and functions of calcium-dependent protein kinases and their relatives in Arabidopsis thaliana. New Phytol. 224, 585–604. 10.1111/nph.16088.

28. Dubiella, U., Seybold, H., Durian, G., Komander, E., Lassig, R., Witte, C.P., Schulze, W.X., and Romeis, T. (2013). Calcium-dependent protein kinase/NADPH oxidase activation circuit is required for rapid defense signal propagation. Proc. Natl. Acad. Sci. U. S. A. 110, 8744–8749. 10.1073/PNAS.1221294110/-/DCSUPPLEMENTAL.

29. Dammann, C., Ichida, A., Hong, B., Romanowsky, S.M., Hrabak, E.M., Harmon, A.C., Pickard, B.G., and Harper, J.F. (2003). Subcellular targeting of nine calcium-dependent protein kinase isoforms from Arabidopsis. Plant Physiol. 132, 1840–1848. 10.1104/PP.103.020008.

30. Lu, S.X., and Hrabak, E.M. (2002). An Arabidopsis Calcium-Dependent Protein Kinase Is Associated with the Endoplasmic Reticulum. Plant Physiol. 128, 1008. 10.1104/PP.010770.

31. Huang, D., Sun, Y., Ma, Z., Ke, M., Cui, Y., Chen, Z., Chen, C., Ji, C., Tran, T.M., Yang, L., et al. (2019). Salicylic acid-mediated plasmodesmal closure via Remorin-dependent lipid organization. Proc. Natl. Acad. Sci. U. S. A. 116, 21274–21284. 10.1073/pnas.1911892116.

32. Mori, I.C., Murata, Y., Yang, Y., Munemasa, S., Wang, Y.F., Andreoli, S., Tiriac, H., Alonso, J.M., Harper, J.F., Ecker, J.R., et al. (2006). CDPKs CPK6 and CPK3 function in ABA regulation of guard cell S-type anion-and Ca2+-permeable channels and stomatal closure. PLoS Biol. 4, 1749–1762. 10.1371/journal.pbio.0040327.

33. Zhang, M., Chang, H., Zhang, Y., Yu, J., Wu, L., Ji, W., Chen, J., Liu, B., Lu, J., Liu, Y., et al. (2012). Rational design of true monomeric and bright photoactivatable fluorescent proteins. Nat. Methods 2012 97 9, 727–729. 10.1038/nmeth.2021.

34. Levet, F., Hosy, E., Kechkar, A., Butler, C., Beghin, A., Choquet, D., and Sibarita, J.B. (2015). SR-Tesseler: a method to segment and quantify localization-based super-resolution microscopy data. Nat. Methods 2015 1211 12, 1065–1071. 10.1038/nmeth.3579.

35. Sheen, J. (1996). Ca2+-dependent protein kinases and stress signal transduction in plants. Science 274, 1900–1902. 10.1126/SCIENCE.274.5294.1900.

36. Harper, J.R., Breton, G., and Harmon, A. (2004). Decoding Ca(2+) signals through plant protein kinases. Annu. Rev. Plant Biol. 55, 263–288. 10.1146/ANNUREV.ARPLANT.55.031903.141627.

37. Huimin, R., Hussain, J., Wenjie, L., Fenyong, Y., Junjun, G., Youhan, K., Shenkui, L., and Guoning, Q. (2021). The expression of constitutively active CPK3 impairs potassium uptake and transport in Arabidopsis under low K + stress. Cell Calcium 98. 10.1016/J.CECA.2021.102447.

38. Sezgin, E., Levental, I., Mayor, S., and Eggeling, C. (2017). The mystery of membrane organization: composition, regulation and roles of lipid rafts. Nat. Rev. Mol. Cell Biol. 18, 361–374. 10.1038/NRM.2017.16.

39. He, J.X., Fujioka, S., Li, T.C., Kang, S.G., Seto, H., Takatsuto, S., Yoshida, S., and Jang, J.C. (2003). Sterols Regulate Development and Gene Expression inArabidopsis. Plant Physiol. 131, 1258. 10.1104/PP.014605.

40. Simon, M.L.A., Platre, M.P., Marquès-Bueno, M.M., Armengot, L., Stanislas, T., Bayle, V., Caillaud, M.C., and Jaillais, Y. (2016). A PI4P-driven electrostatic field controls cell membrane identity and signaling in plants. Nat. plants 2, 16089. 10.1038/NPLANTS.2016.89.

41. Raffaele, S., Mongrand, S., Gamas, P., Niebel, A., and Ott, T. (2007). Genome-wide annotation of remorins, a plant-specific protein family: Evolutionary and functional perspectives. Plant Physiol. 145, 593–600. 10.1104/pp.107.108639.

42. Mergner, J., Frejno, M., List, M., Papacek, M., Chen, X., Chaudhary, A., Samaras, P., Richter, S., Shikata, H., Messerer, M., et al. (2020). Mass-spectrometry-based draft of the Arabidopsis proteome. Nature 579, 409–414. 10.1038/S41586-020-2094-2.

43. Lefebvre, B., Timmers, T., Mbengue, M., Moreau, S., Hervé, C., Tóth, K., Bittencourt-Silvestre, J., Klaus, D., Deslandes, L., Godiard, L., et al. (2010). A remorin protein interacts with symbiotic receptors and regulates bacterial infection. Proc. Natl. Acad. Sci. U. S. A. 107, 2343–2348. 10.1073/pnas.0913320107.

44. Gouguet, P., Gronnier, J., Legrand, A., Perraki, A., Jolivet, M.D., Deroubaix, A.F., Retana, S.G., Boudsocq, M., Habenstein, B., Mongrand, S., et al. (2021). Connecting the dots: From nanodomains to physiological functions of REMORINs. Plant Physiol. 185, 632–649. 10.1093/PLPHYS/KIAA063.

45. Abel, N.B., Buschle, C.A., Hernandez-Ryes, C., Burkart, S.S., Deroubaix, A.-F., Mergner, J., Gronnier, J., Jarsch, I.K., Folgmann, J., Braun, K.H., et al. (2021). A hetero-oligomeric remorin-receptor complex regulates plant development. bioRxiv, 2021.01.28.428596. 10.1101/2021.01.28.428596.

46. Wang, Y., Gong, Q., Wu, Y., Huang, F., Ismayil, A., Zhang, D., Li, H., Gu, H., Ludman, M., Fátyol, K., et al. (2021). A calmodulin-binding transcription factor links calcium signaling to antiviral RNAi defense in plants. Cell Host Microbe 29, 1393–1406.e7. 10.1016/j.chom.2021.07.003.

47. Dong, C.H., and Hong, Y. (2013). Arabidopsis CDPK6 phosphorylates ADF1 at N-terminal serine 6 predominantly. Plant Cell Rep. 32, 1715–1728. 10.1007/S00299-013-1482-6.

48. Wang, J., Lian, N., Zhang, Y., Man, Y., Chen, L., Yang, H., Lin, J., and Jing, Y. (2022). The Cytoskeleton in Plant Immunity: Dynamics, Regulation, and Function. Int. J. Mol. Sci. 2022, Vol. 23, Page 15553 23, 15553. 10.3390/IJMS232415553.

49. Harries, P.A., Park, J.-W., Sasaki, N., Ballard, K.D., Maule, A.J., and Nelson, R.S. Differing requirements for actin and myosin by plant viruses for sustained intercellular movement.

50. Tilsner, J., Linnik, O., Wright, K.M., Bell, K., Roberts, A.G., Lacomme, C., Cruz, S.S., and Oparka, K.J. (2012). The TGB1 movement protein of potato virus X reorganizes actin and endomembranes into the X-body, a viral replication factory. Plant Physiol. 158, 1359–1370. 10.1104/pp.111.189605.

51. Köster, P., DeFalco, T.A., and Zipfel, C. (2022). Ca2+ signals in plant immunity. EMBO J. 41, e110741. 10.15252/EMBJ.2022110741.

52. Saito, S., Hamamoto, S., Moriya, K., Matsuura, A., Sato, Y., Muto, J., Noguchi, H., Yamauchi, S., Tozawa, Y., Ueda, M., et al. (2018). N-myristoylation and S-acylation are common modifications of Ca2+-regulated Arabidopsis kinases and are required for activation of the SLAC1 anion channel. New Phytol. 218, 1504–1521. 10.1111/nph.15053.

53. Asai, S., Ichikawa, T., Nomura, H., Kobayashi, M., Kamiyoshihara, Y., Mori, H., Kadota, Y., Zipfel, C., Jones, J.D.G., and Yoshioka, H. (2013). The variable domain of a plant calcium-dependent protein kinase (CDPK) confers subcellular localization and substrate recognition for NADPH oxidase. J. Biol. Chem. 288, 14332–14340. 10.1074/jbc.M112.448910.

54. Valmonte-Cortes, G.R., Lilly, S.T., Pearson, M.N., Higgins, C.M., and Macdiarmid, R.M. (2022). The Potential of Molecular Indicators of Plant Virus Infection: Are Plants Able to Tell Us They Are Infected? Plants 11, 188. 10.3390/PLANTS11020188/S1.

55. Su, C., Rodriguez-Franco, M., Lace, B., Nebel, N., Hernandez-Reyes, C., Liang, P., Schulze, E., Mymrikov, E. V., Gross, N.M., Knerr, J., et al. (2023). Stabilization of membrane topologies by proteinaceous remorin scaffolds. Nat. Commun. 2023 141 14, 1–16. 10.1038/s41467-023-35976-5.

56. Perraki, A., Cacas, J.L., Crowet, J.M., Lins, L., Castroviejo, M., German-Retana, S., Mongrand, S., and Raffaele, S. (2012). Plasma membrane localization of Solanum tuberosum Remorin from group 1, homolog 3 is mediated by conformational changes in a novel C-terminal anchor and required for the restriction of potato virus X movement. Plant Physiol. 160, 624–637. 10.1104/pp.112.200519.

57. Gronnier, J., Crowet, J.-M., Habenstein, B., Nasir, M.N., Bayle, V., Hosy, E., Platre, M.P., Gouguet, P., Raffaele, S., Martinez, D., et al. (2017). Structural basis for plant plasma membrane protein dynamics and organization into functional nanodomains. Elife 6. 10.7554/eLife.26404.

58. Konrad, S.S.A., Popp, C., Stratil, T.F., Jarsch, I.K., Thallmair, V., Folgmann, J., Marín, M., and Ott, T. (2014). S-acylation anchors remorin proteins to the plasma membrane but does not primarily determine their localization in membrane microdomains. New Phytol. 203, 758–769. 10.1111/nph.12867.

59. Smokvarska, M., Jaillais, Y., and Martiniere, A. (2021). Function of membrane domains in rho-of-plant signaling. Plant Physiol. 185, 663–681. 10.1093/PLPHYS/KIAA082.

60. Tran, T.M., Ma, Z., Triebl, A., Nath, S., Cheng, Y., Gong, B.Q., Han, X., Wang, J., Li, J.F., Wenk, M.R., et al. (2020). The bacterial quorum sensing signal DSF hijacks Arabidopsis thaliana sterol biosynthesis to suppress plant innate immunity. Life Sci. Alliance 3. 10.26508/LSA.202000720.

61. Su, C., Klein, M.-L., Hernández-Reyes, C., Batzenschlager, M., Ditengou, F.A., Lace, B., Keller, J., Delaux, P.-M., and Ott, T. (2020). The Medicago truncatula DREPP Protein Triggers Microtubule Fragmentation in Membrane Nanodomains during Symbiotic Infections. Plant Cell, tpc.00777.2019. 10.1105/tpc.19.00777.

62. Grison, M.S., Kirk, P., Brault, M.L., Wu, X.N., Schulze, W.X., Benitez-Alfonso, Y., Immel, F., and Bayer, E.M. (2019). Plasma membrane-associated receptor-like kinases relocalize to plasmodesmata in response to osmotic stress. Plant Physiol. 181, 142–160. 10.1104/pp.19.00473.

63. Jarsch, I.K., and Ott, T. (2011). Perspectives on Remorin Proteins, Membrane Rafts, and Their Role During Plant–Microbe Interactions. MPMI 19. 10.1094/MPMI.

64. Gui, J., Zheng, S., Liu, C., Shen, J., Li, J., and Li, L. (2016). OsREM4.1 Interacts with OsSERK1 to Coordinate the Interlinking between Abscisic Acid and Brassinosteroid Signaling in Rice. Dev. Cell 38, 201–213. 10.1016/j.devcel.2016.06.011.

65. Teixeira, R.M., Ferreira, M.A., Raimundo, G.A.S., Loriato, V.A.P., Reis, P.A.B., and Fontes, E.P.B. (2019). Virus perception at the cell surface: revisiting the roles of receptor-like kinases as viral pattern recognition receptors. Mol. Plant Pathol. 20, 1196–1202. 10.1111/MPP.12816.

66. Liang, P., Stratil, T.F., Popp, C., Marín, M., Folgmann, J., Mysore, K.S., Wen, J., and Ott, T. (2018). Symbiotic root infections in Medicago truncatula require remorin-mediated receptor stabilization in membrane nanodomains. Proc. Natl. Acad. Sci. U. S. A. 115, 5289–5294. 10.1073/pnas.1721868115.

67. Gronnier, J., Franck, C.M., Stegmann, M., Defalco, T.A., Abarca, A., von Arx, M., Dünser, K., Lin, W., Yang, Z., Kleine-Vehn, J., et al. (2022). Regulation of immune receptor kinase plasma membrane nanoscale organization by a plant peptide hormone and its receptors. Elife 11. 10.7554/ELIFE.74162.

68. Szymanski, W.G., Zauber, H., Erban, A., Gorka, M., Wu, X.N., and Schulze, W.X. (2015). Cytoskeletal components define protein location to membrane microdomains. Mol. Cell. Proteomics 14, 2493–2509. 10.1074/mcp.M114.046904.

69. Stegmann, M., Monaghan, J., Smakowska-Luzan, E., Rovenich, H., Lehner, A., Holton, N., Belkhadir, Y., and Zipfel, C. (2017). The receptor kinase FER is a RALF-regulated scaffold controlling plant immune signaling. Science (80-. ). 355, 287–289. 10.1126/SCIENCE.AAL2541/SUPPL_FILE/STEGMANN_SM_REVISION3.PDF.

70. Noack, L.C., Bayle, V., Armengot, L., Rozier, F., Mamode-Cassim, A., Stevens, F.D., Caillaud, M.C., Munnik, T., Mongrand, S., Pleskot, R., et al. (2022). A nanodomain-anchored scaffolding complex is required for the function and localization of phosphatidylinositol 4-kinase alpha in plants. Plant Cell 34, 302. 10.1093/PLCELL/KOAB135.

71. Boudsocq, M., Droillard, M.J., Regad, L., and Laurière, C. (2012). Characterization of Arabidopsis calcium-dependent protein kinases: Activated or not by calcium? Biochem. J. 447, 291–299. 10.1042/BJ20112072.

72. Concordet, J.P., and Haeussler, M. (2018). CRISPOR: intuitive guide selection for CRISPR/Cas9 genome editing experiments and screens. Nucleic Acids Res. 46, W242–W245. 10.1093/NAR/GKY354.

73. Wang, Z.P., Xing, H.L., Dong, L., Zhang, H.Y., Han, C.Y., Wang, X.C., and Chen, Q.J. (2015). Egg cell-specific promoter-controlled CRISPR/Cas9 efficiently generates homozygous mutants for multiple target genes in Arabidopsis in a single generation. Genome Biol. 16. 10.1186/S13059-015-0715-0.

74. Jarsch, I.K., Konrad, S.S.A., Stratil, T.F., Urbanus, S.L., Szymanski, W., Braun, P., Braun, K.H., and Ott, T. (2014). Plasma membranes are Subcompartmentalized into a plethora of coexisting and diverse microdomains in Arabidopsis and Nicotiana benthamiana. Plant Cell 26, 1698–1711. 10.1105/tpc.114.124446.

75. Rohr L, Ehinger A, Rausch L, Glöckner Burmeister N, Meixner AJ, Gronnier J, Harter K, Kemmerling B and zur Oven-Krockhaus S (2024) OneFlowTraX: a user-friendly software for super-resolution analysis of single-molecule dynamics and nanoscale organization. Front. Plant Sci. 15:1358935. doi: 10.3389/fpls.2024.1358935

