## Supplementary material for "Interdependence of plasma membrane nanoscale dynamics of a kinase and its cognate substrate underlies *Arabidopsis* response to viral infection": Figure 4 Supplemental Figure 3

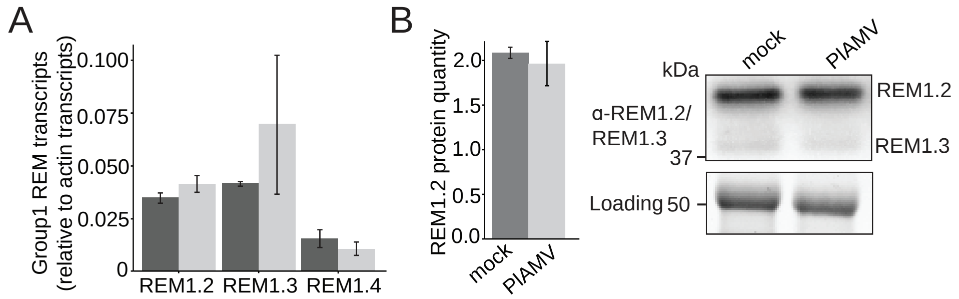


Figure 4 - figure supplement 3. Group 1 REMs transcript and protein levels are not modified during a PlAMV infection
