## Supplementary figures and images for "Interdependence of plasma membrane nanoscale dynamics of a kinase and its cognate substrate underlies *Arabidopsis* response to viral infection"

### Figure 1 Supplemental Figure 1

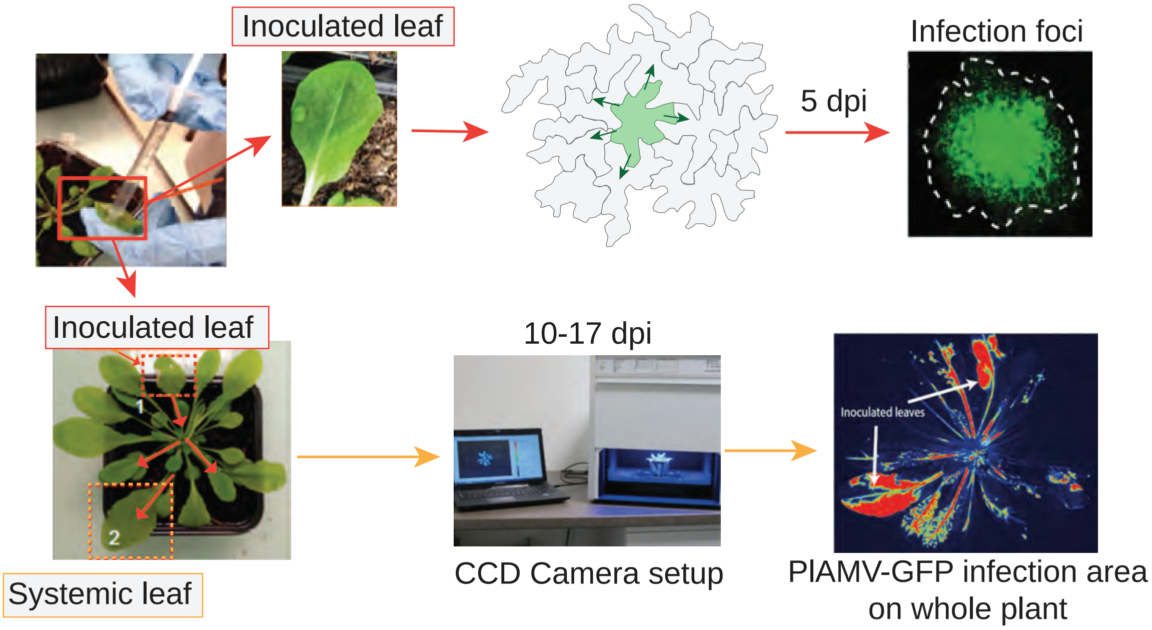


Figure 1 - figure supplement 1. Viral propagation experimental design

### Figure 1 Supplemental Figure 2

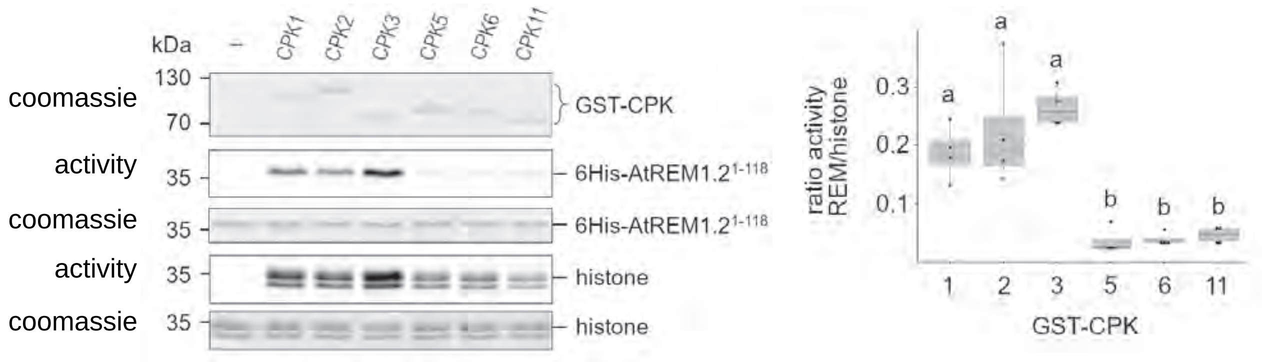


Figure 1 - figure supplement 2. Specificity of CPK kinase activity towards REM1.2 *in vitro*

### Figure 1 Supplemental Figure 3

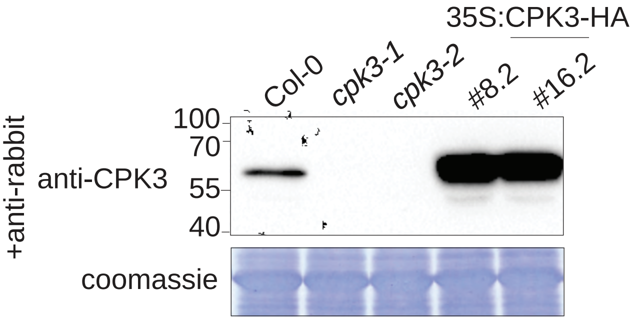


Figure 1 - figure supplement 3. CPK3 protein levels in knock-out and overexpressing lines

### Figure 2 Supplemental Figure 2

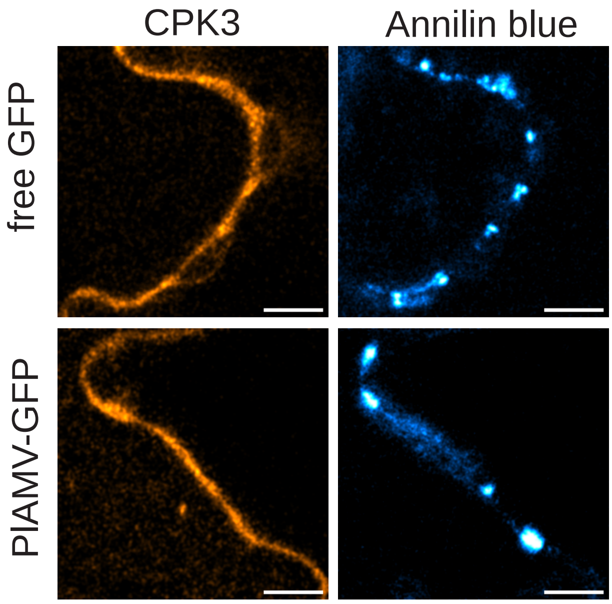


Figure 2 – figure supplement 2. CPK3 does not accumulate at plasmodesmata during a PlAMV infection

### Figure 3 Supplemental Figure 1

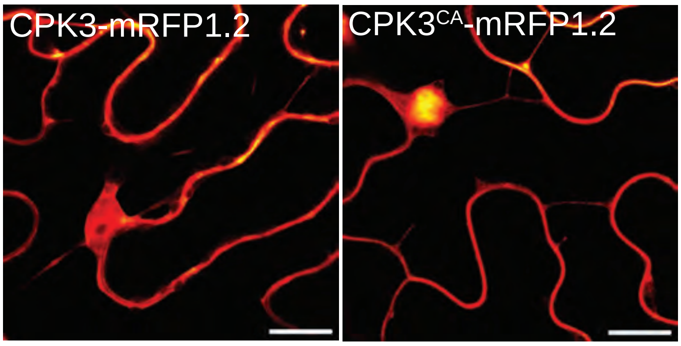


Figure 3 - figure supplement 1. CPK3 and CPK3$^{CA}$ display a similar subcellular localization

### Figure 4 Supplemental Figure 1

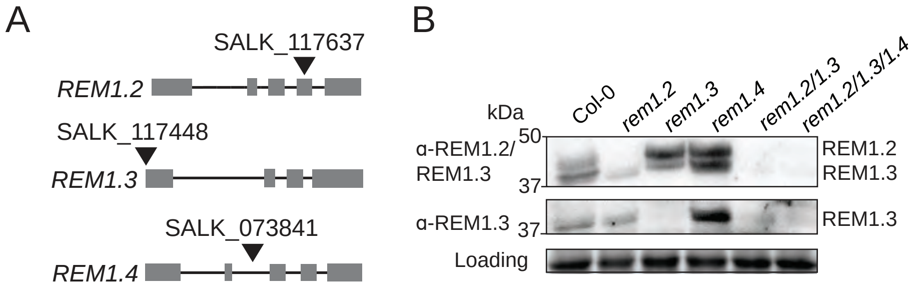


Figure 4 - figure supplement 1. Group 1 REM single, double and triple knock out mutants

### Figure 4 Supplemental Figure 4

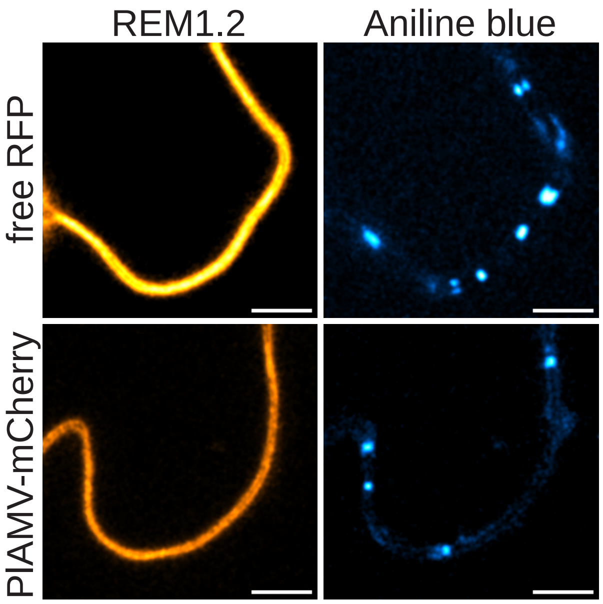


Figure 4 – figure supplement 4. REM1.2 does not accumulate at plasmodesmata during a PlAMV infection

### Figure 5 Supplemental Figure 1

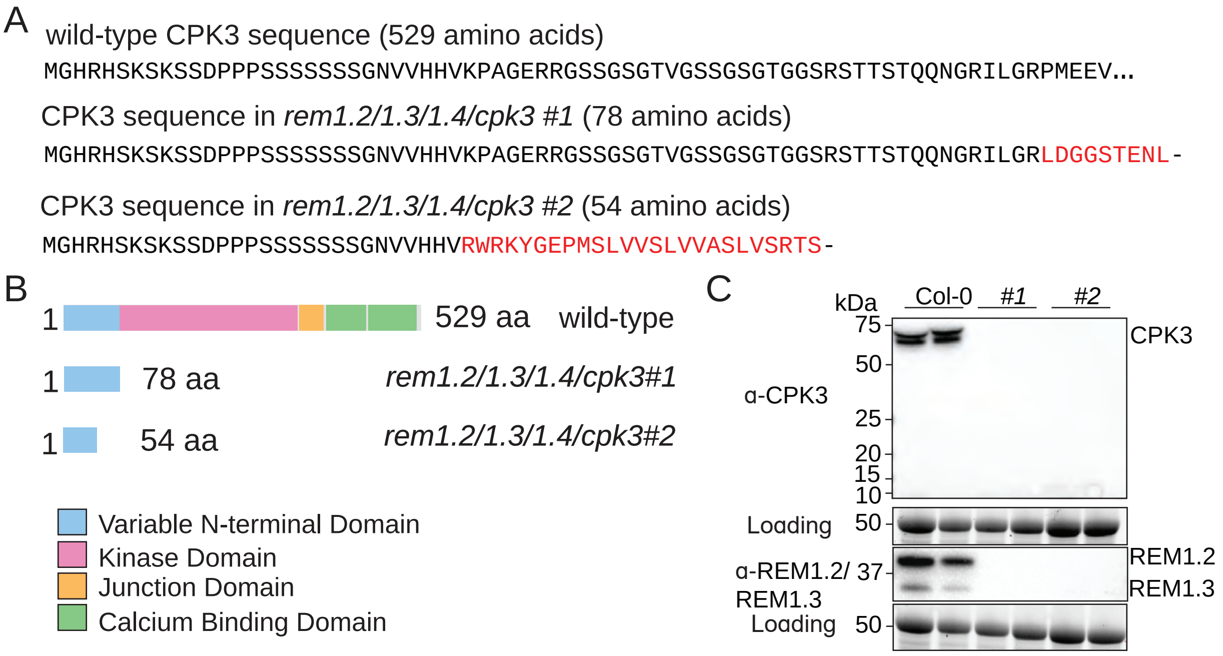


Figure 5 - figure supplement 1. CRISPR-mediated knock out of cpk3 in *rem1.2 rem1.3 rem1.4* background
