## Supplementary material for "Interdependence of plasma membrane nanoscale dynamics of a kinase and its cognate substrate underlies *Arabidopsis* response to viral infection": Figure 5 Supplemental Figure 2

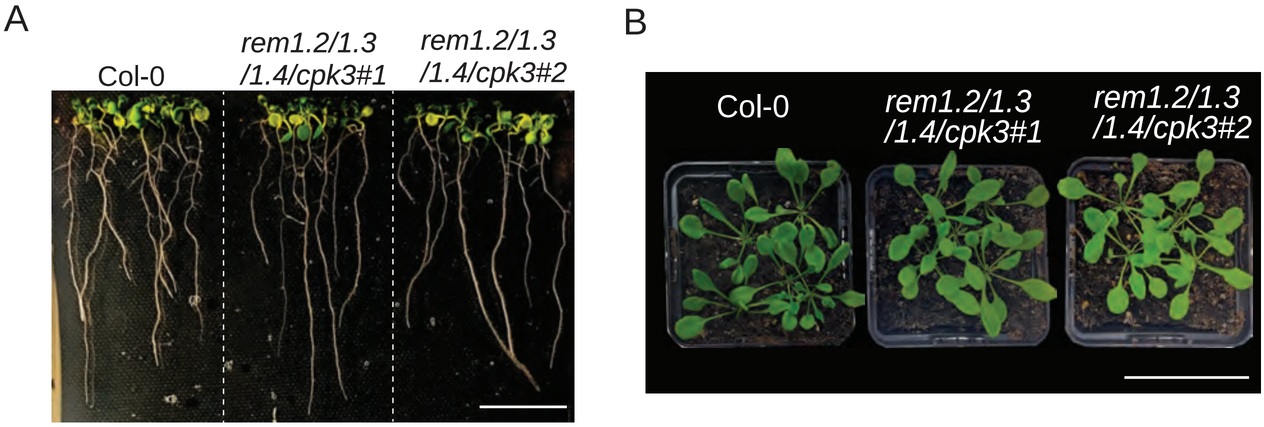


Figure 5 - figure supplement 2. *rem1.2 rem1.3 rem1.4 cpk3* quadruple knock-out lines do not display any obvious developmental phenotype
