## Supplementary material for "Interdependence of plasma membrane nanoscale dynamics of a kinase and its cognate substrate underlies *Arabidopsis* response to viral infection": Figure 5 Supplemental Figure 3

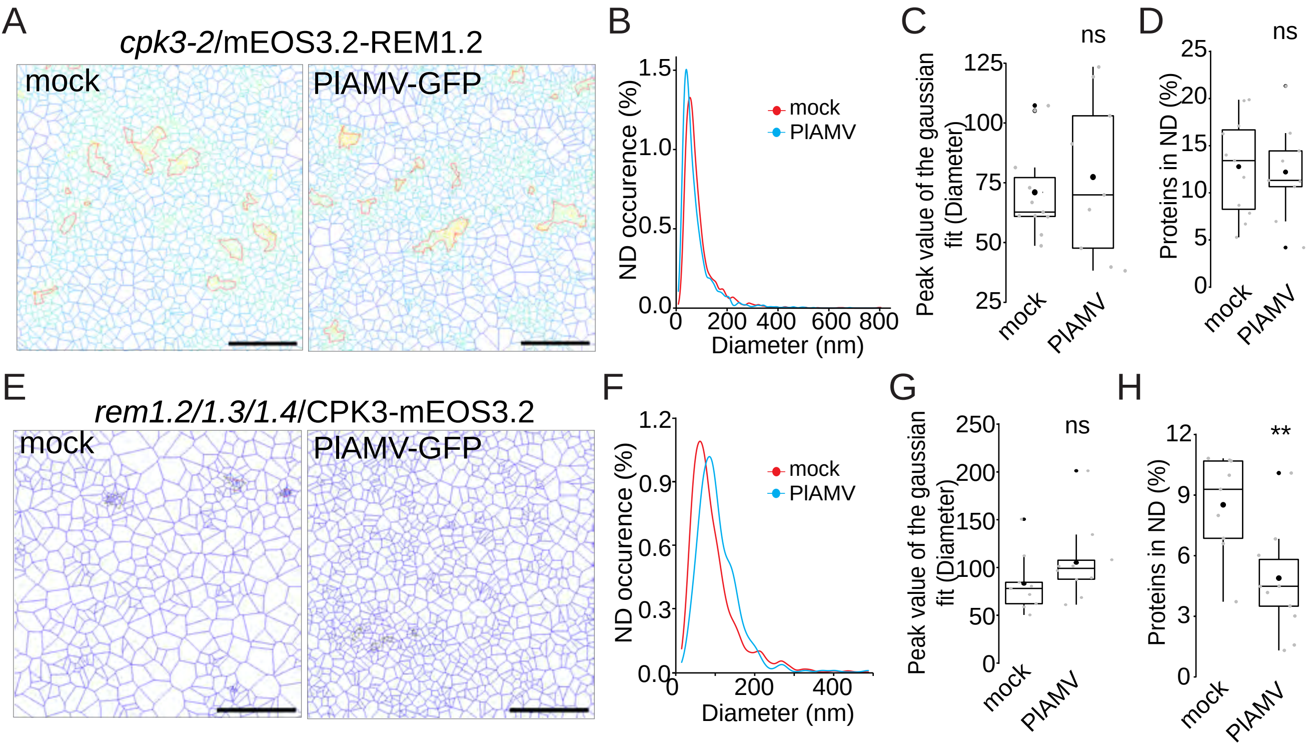


Figure 5 - figure supplement 3. CPK3 and group1 REM role in each other PM nanoorganization during PlAMV infection
