## Supplementary material for "Interdependence of plasma membrane nanoscale dynamics of a kinase and its cognate substrate underlies *Arabidopsis* response to viral infection": Figure 5 Supplemental Figure 4

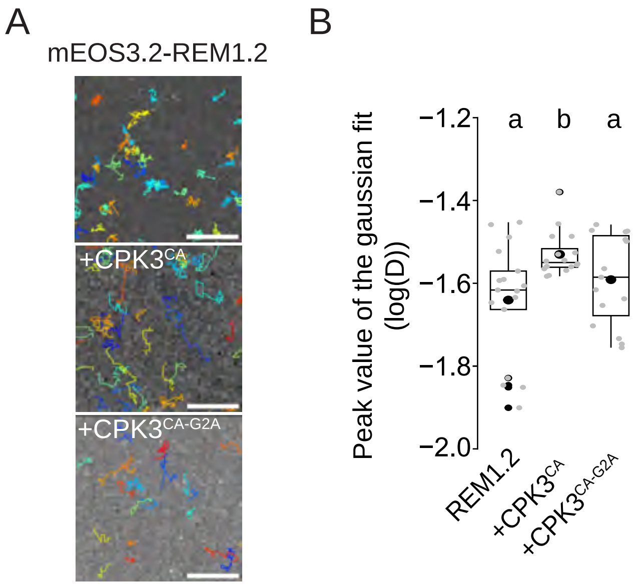


Figure 5 - figure supplement 4. REM1.2 diffusion is increased upon co-expression with CPK3$^{CA}$ but not with CPK3$^{CA\text{-}G2A}$
