## Supplementary material for "Interdependence of plasma membrane nanoscale dynamics of a kinase and its cognate substrate underlies *Arabidopsis* response to viral infection": Figure 5 Supplemental Figure 5

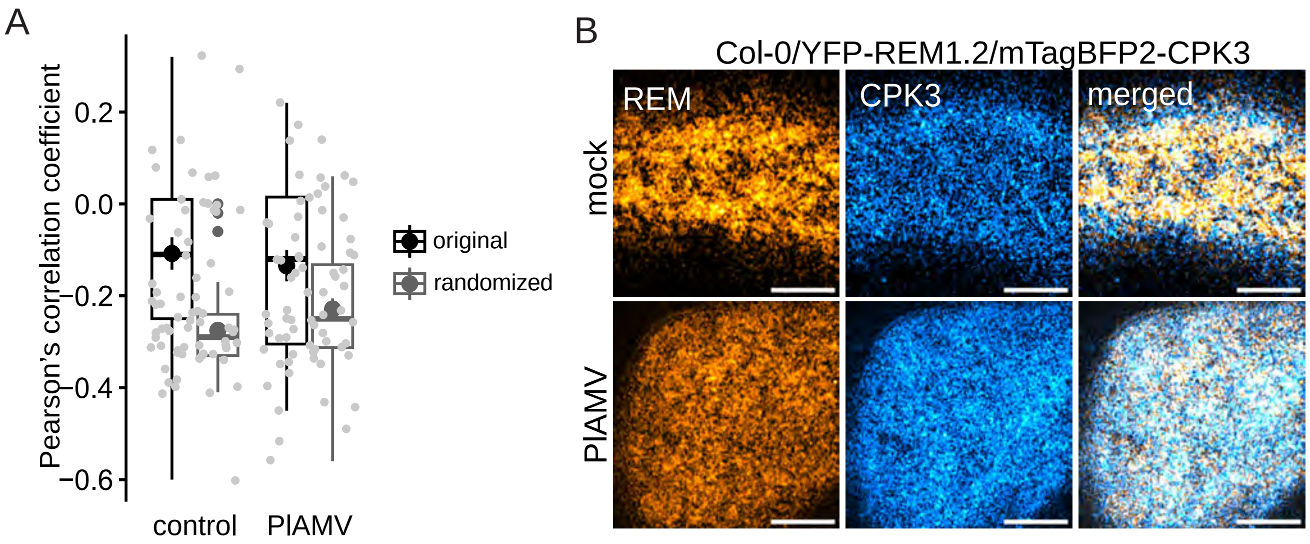


Figure 5 - figure supplement 5. PlAMV infection does not modify CPK3 and group1 REM colocalization coefficient
