## Supplementary material for "Interdependence of plasma membrane nanoscale dynamics of a kinase and its cognate substrate underlies *Arabidopsis* response to viral infection": Figure 2 Supplemental Figure 1

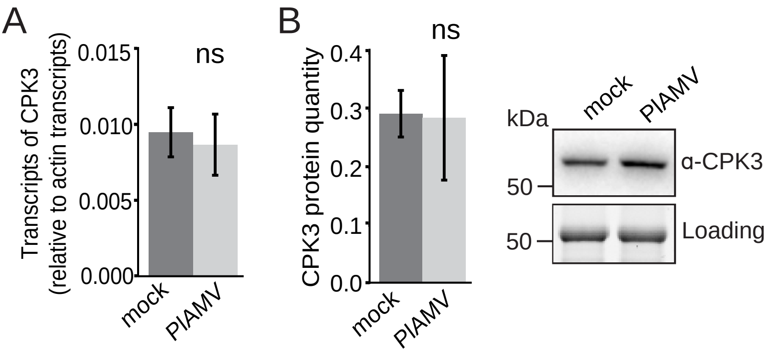


Figure 2 – figure supplement 1. CPK3 transcript and proteins levels are not modified upon viral infection
