## Supplementary material for "Interdependence of plasma membrane nanoscale dynamics of a kinase and its cognate substrate underlies *Arabidopsis* response to viral infection": Figure 4 Supplemental Figure 2

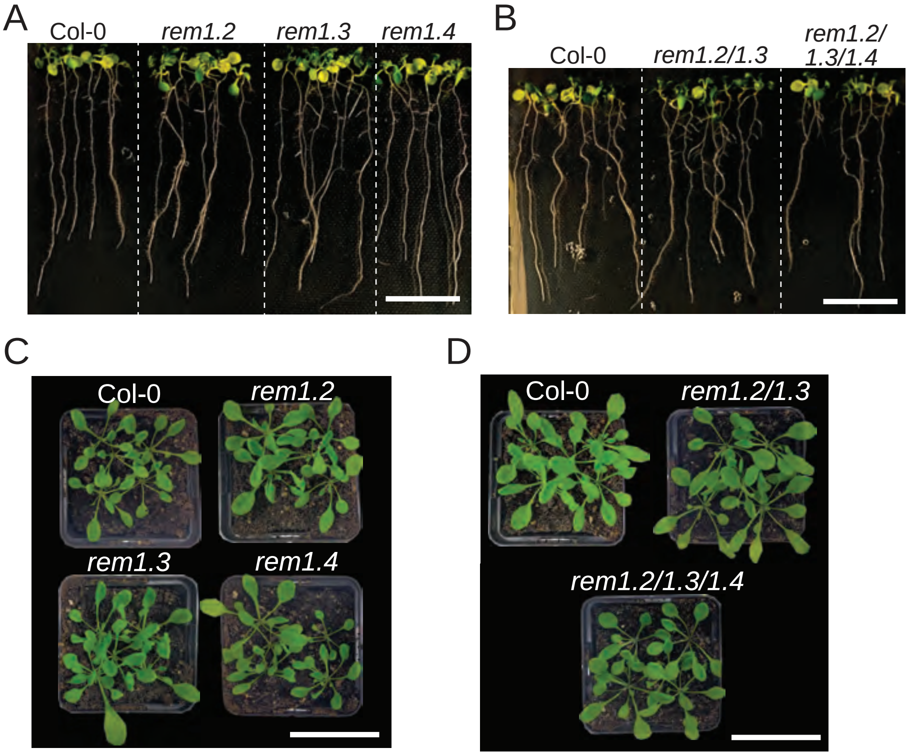


Figure 4 - figure supplement 2. Group 1 REMs single and multiple knock out lines do not display any obvious developmental phenotype
